# SMAD4 target genes are part of a transcriptional network that integrates the response to BMP and SHH signaling during early limb bud patterning

**DOI:** 10.1101/2021.09.08.459466

**Authors:** Julie Gamart, Iros Barozzi, Frédéric Laurent, Robert Reinhardt, Laurène Ramos Martins, Thomas Oberholzer, Axel Visel, Rolf Zeller, Aimée Zuniga

## Abstract

SMAD4 regulates gene expression in response to BMP and TGFβ signal transduction and is required for diverse morphogenetic processes, but its target genes have remained largely elusive. Here, we use an epitope-tagged *Smad4* allele for ChIP-seq analysis together with transcriptome analysis of wild-type and mouse forelimb buds lacking *Smad4* in the mesenchyme. This analysis identifies the SMAD4 target genes during establishment of the feedback signaling system and establishes that SMAD4 predominantly mediates BMP signal-transduction during early limb bud development. Unexpectedly, the initial analysis reveals that the expression of cholesterol biosynthesis enzymes is precociously down-regulated and intracellular cholesterol levels reduced in *Smad4*-deficient limb bud mesenchymal progenitors. The SMAD4 target GRNs includes genes, whose expression in the anterior limb bud is up-regulated by interactions of SMAD4 complexes with enhancers active in the anterior mesenchyme. This reveals a predominant function of SMAD4 in up-regulating target gene expression in the anterior limb bud mesenchyme. Analysis of differentially expressed genes that are shared between *Smad4*- and *Shh*-deficient limb buds corroborates the positive role of SMAD4 in transcriptional regulation of anterior genes and reveals a repressive effect on posterior genes that are positively regulated by SHH signaling. This analysis uncovers the overall opposing effects of SMAD4-mediated BMP and SHH signalling on transcriptional regulation during early limb bud development. In summary, this analysis indicates that during early digit patterning and limb bud outgrowth, the anterior/proximal and proximo/distal expression dynamics of co-regulated genes are controlled by distinct and contrasting trans-regulatory inputs from SHH and SMAD4-mediated BMP signal transduction.

## INTRODUCTION

The transforming growth factor-beta (TGFβ) and bone morphogenetic protein (BMP) pathway constitutes one of the major signaling pathways that control vertebrate embryonic development (reviewed in Weiss and Attisano, 2013). Of relevance to the present study, BMPs are required for limb bud formation and outgrowth (reviewed in Pignatti et al., 2014). BMP ligands activate their transmembrane BMP receptors (BMPR1A/1B isoforms and BMPR2) that form hetero-tetrameric complexes. The activated BMP receptor complexes trigger R-SMAD (SMAD1,-5,-8) phosphorylation; R-SMADs in turn form a complex with SMAD4, which translocates to the nucleus to regulate target gene expression by recruiting either co-activators or repressors. SMAD4 is part of the SMAD complexes that bind to DNA in response to TGFβ/BMP signal transduction (Weiss and Attisano, 2013). Genetic studies in mice have identified several functions of BMP signaling during limb bud development (Pignatti et al., 2014). Limb bud initiation and early outgrowth (mouse embryonic days E9.5-E10.0) require high mesenchymal BMP4 activity together with BMPR1A and SMAD4-mediated signal transduction in the ectoderm to establish the apical ectodermal ridge (AER) as the fibroblast growth factor (FGF) signaling centre (Ahn et al., 2001; Bénazet and Zeller, 2013; Pajni-Underwood et al., 2007; see also Pignatti et al., 2014). Genetic inactivation of *Bmp4* during forelimb bud formation completely disrupts outgrowth and initiation of the expression of the BMP antagonist *Gremlin1* (*Grem1*; Bénazet et al., 2009). In turn, BMP-mediated activation and up-regulation of *Grem1* expression progressively lowers mesenchymal BMP activity, which is reinforced by Sonic Hedgehog (SHH) signaling as part of the self-regulatory SHH/GREM1/AER-FGF feedback signaling system (*Grem1*; Bénazet et al., 2009). Genetic analysis in the mouse has shown that *Shh* is transiently required during early limb bud development to specify digit identities and subsequently to promote the proliferative expansion of the limb bud mesenchymal progenitors (LMPs) that give rise to the digit primordia (Zhu et al., 2008). Furthermore, genetic inactivation of *Grem1* disrupts the self-regulatory signaling system and thereby digit patterning and distal limb bud development, which is restored by lowering *Bmp4* genetically in *Grem1*-deficient mouse embryos (Bénazet et al., 2009). These results show that limb bud patterning and outgrowth depend critically on GREM1-mediated reduction of BMP activity, morphogenetic SHH signaling and proliferation of LMPs as part of the SHH/GREM1/AER-FGF signaling system (Bénazet et al., 2009; Probst et al., 2011). In addition, mesenchymal BMP activity is essential to regulate AER length, which in turn prevents digit polydactyly (Bénazet et al., 2009; Lopez-Rios et al., 2012; Selever et al., 2004). During limb bud outgrowth, SMAD4-mediated signal transduction in the mesenchyme is required together with SHH signaling for positive regulation and propagation of *Grem1* expression (Bénazet et al., 2012). During handplate formation, mesenchymal SMAD4 is required for termination of *Grem1* expression and the self-regulatory SHH/GREM1/AER-FGF signaling system (Scherz et al., 2004; Verheyden and Sun, 2008), as both Shh and AER-*Fgf8* expression are prolonged in mouse limb buds lacking *Smad4* (Bénazet et al., 2012). In addition, the genetic analysis showed that SMAD4-mediated BMP signal transduction is required to initiate the aggregation and differentiation of the chondrogenic progenitors giving rise to the cartilage primordia of all limb skeletal elements (Bénazet et al., 2012; Lopez-Rios et al., 2012; Pizette and Niswander, 2000). In contrast, there is no genetic data pointing to essential functions of TGFβ signaling during the early phase of limb bud outgrowth and patterning that is controlled by the self-regulatory signaling system. However, a study using limb bud cell cultures has provided evidence that TGFβ signaling alleviates an inhibitory effect of BMPs in specification of *Sox9*-positive osteochondrogenic progenitors (Karamboulas et al., 2010).

To gain an unbiased view of how SMAD4 mediates signal transduction during early mouse forelimb bud development, we have identified its direct transcriptional targets (i.e. SMAD4 target genes) using a novel *Smad4* allele with an inserted 3XFLAG epitope tag (*Smad4*^3xF^ allele). This *Smad4*^3xF^ allele allows for sensitive and unbiased detection of the genomic regions enriched in endogenous SMAD4-chromatin complexes using ChIP-seq analysis. Combining the SMAD4 cistrome with RNA-seq analysis of wild-type and *Smad4*^Δ/Δc^ mouse forelimb buds (lacking *Smad4* in the mesenchyme) identifies the SMAD4 target genes among the differentially expressed genes (DEGs) during the onset (E10.0) and early progression (E10.5) of limb bud outgrowth and patterning. This analysis reveals that one unexpected SMAD4 function is its requirement for maintaining the expression of the majority of the cholesterol biosynthesis enzymes in early limb buds (E10.0). In *Smad4*^Δ/Δc^ forelimb buds their expression is precociously down-regulated, which is paralleled by a reduction of endogenous cholesterol levels in mutant LMPs. We also identify the direct SMAD4 targets in the TGFß and BMP pathways, which establishes that in early forelimb buds (at least up to E10.5) the *Smad4* deficiency preferentially disrupts BMP signal-transduction. A whole mount *in situ* hybridisation screen identifies SMAD4 target genes whose spatial expression is altered in early limb buds. Furthermore, *LacZ reporter* analysis shows that the associated enhancers for some of these target genes that are enriched in SMAD4-chromatin complexes are active preferentially in the anterior forelimb bud mesenchyme. Together with the observed spatial changes in gene expression, this points to SMAD4 functions in upregulating target gene expression in the anterior limb bud mesenchyme. Comparative analysis of the DEGs in *Smad4*^Δ/Δc^ and *Shh*^Δ/Δc^ forelimb buds (Probst et al., 2011) identifies the genes co-regulated by both pathways in early limb buds. Gene regulatory network (GRN) analysis reveals the interactions of SMAD4-mediated BMP signal transduction with SHH signaling in regulating the spatial expression of key regulator genes during the early transient phase of digit specification (Zhu et al., 2008) and establishment of self-regulatory limb bud signaling.

## RESULTS

### Identification of the SMAD4 cistrome and target genes in mouse forelimb buds

Specific detection of the endogenous SMAD4 protein complexes was achieved by inserting a 3xFLAG (3xF) epitope tag into the SMAD4 carboxy-terminal domain using homologous recombination in mouse ES cells (Fig. S1). This does not alter *Smad4* transcript levels and homozygous *Smad4*^3xF/3xF^ embryos are born at the expected Mendelian ratios. In particular, insertion of the 3xFLAG epitope tag does not affect SMAD4 functions during limb bud development as no alterations in chondrogenesis and skeletal development are observed in contrast to mouse limb buds lacking mesenchymal *Smad4* (Bénazet et al., 2012). In early mouse forelimb buds, the distribution of SMAD4^3xF^ proteins is uniform. Higher levels of SMAD4 proteins are detected in the cytoplasm than in the nucleus, but diffuse nucleoplasmic staining is detected in most mesenchymal cells of *Smad4*^3xF/3xF^ forelimb buds at E9.5 (Figure S1, see also Bénazet et al., 2012).

Forelimb buds of *Smad4*^3xF/3xF^ embryos at two stages were used to identify the SMAD4 cistromes during the onset of limb bud development with high mesenchymal BMP activity (E9.5-E10.0, 25-30 somites) and during early outgrowth, when BMP signal transduction is lowered by GREM1-mediated BMP antagonism (E10.5, 34-38 somites, Bénazet et al., 2009) The SMAD4 cistrome of forelimb buds at both stages was determined using chromatin immunoprecipitation in combination with next generation sequencing (ChIP-seq, Fig. 1). Two biological replicates consisting each of ∼80 dissected forelimb buds were analysed per stage. The dissected forelimb buds included some proximal trunk tissue to also detect interactions with genes expressed early in the proximal limb bud and flank mesenchyme. For early limb buds (E9.5-E10.0), statistical analysis of the two replicates by MACS and MSPC identified 2,073 significantly and reproducibly enriched SMAD4 ChIP-seq peaks (Jalili et al., 2015; see materials and methods). About 40% of them are located close to transcriptional start sites (TSS, ±5kb) while ∼20% are located ≥ kb away from TSS (left panel, Fig. 1A). During limb bud outgrowth (E10.5), 6,185 significantly enriched and conserved SMAD4 ChIP-seq peaks were identified, most of which are also located close to TSS (right panel, Fig. 1A). Evolutionary conservation analysis shows that the peak summits of the genomic regions enriched in SMAD4-chromatin complexes are more conserved than the flanking regions in placental mammals (Fig. 1B). Enrichment analyses for known and *de novo* motifs using HOMER identified the SMAD consensus binding motifs as the most enriched motifs at both stages (Figure 1C, 1D; Heinz et al., 2010). In addition, the PKNOX1/PREP1 homeobox motif is the most enriched *de novo* motif in the SMAD4^3xF^ forelimb bud cistrome at E10.5 (right panel, Fig. 1D). This could be functionally relevant as the TALE homeodomain transcription factors PKNOX1/PREP1 and PBX1 interact with SMAD4 to regulate gene expression in cultured cells (Bailey et al., 2004). However, our analysis also revealed significant differences in the overall binding motifs enriched in SMAD4-chromatin complexes from forelimb buds characterized by high (E9.5-E10.0) and low mesenchymal BMP activity (E10.5, Fig. 1C, 1D). Furthermore, the BMP responsive elements (BREs) located near the *Id1* and *Msx2* genes are significantly enriched in the SMAD4^3xF^ ChIP-seq datasets at both stages, which was confirmed by ChIP-qPCR analysis (Fig. 1E). Therefore, the two SMAD4^3xF^ cistromes constitute valid resources to identify the limb bud mesenchymal SMAD4 target genes during the onset (E9.5-E10.0) and distal progression (E10.5) of forelimb bud development.

**Figure 1.**
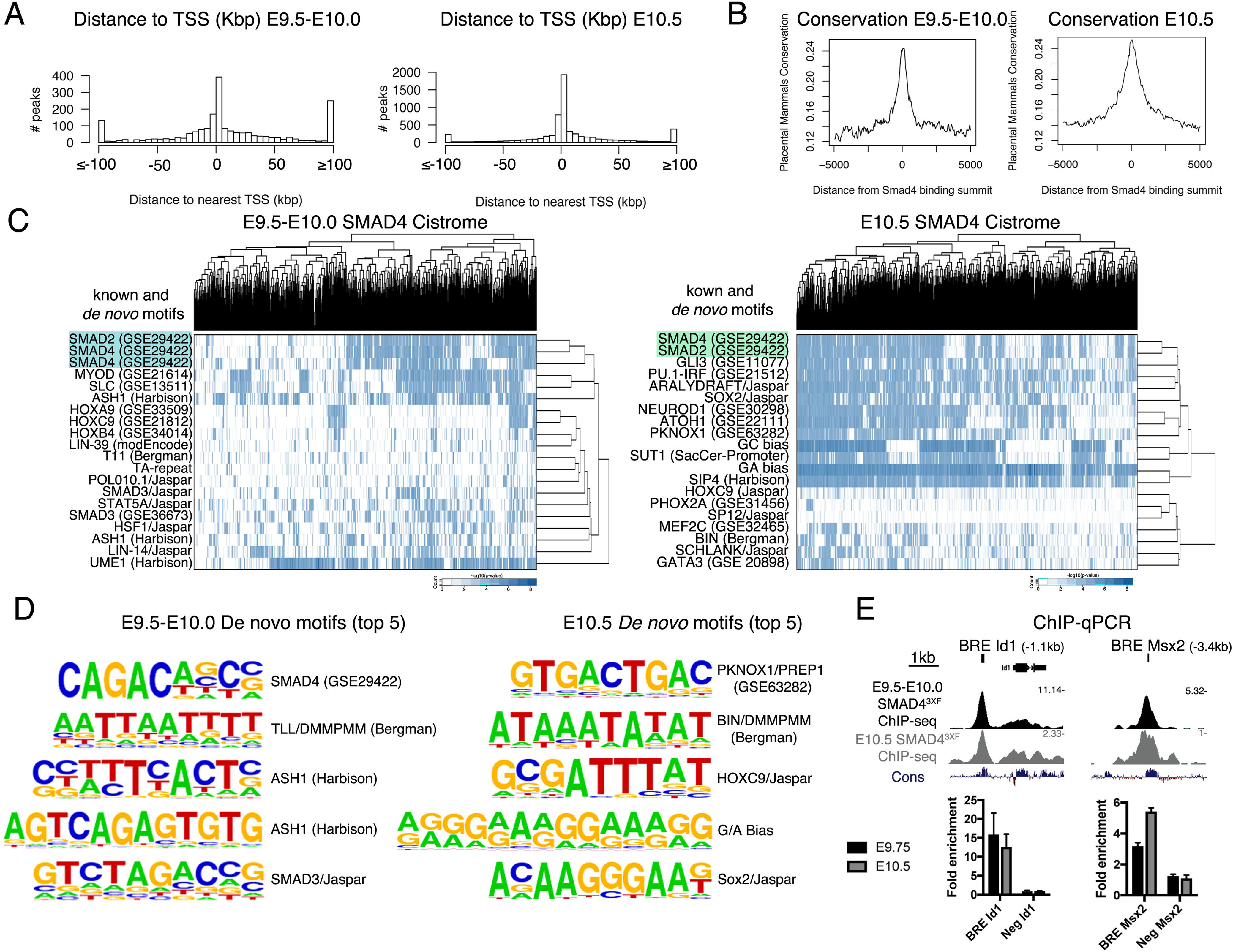
ChIP-seq using the *Smad4^3xF^* allele identifies the genomic regions enriched in SMAD4-chromatin complexes in forelimb buds. (A) Histogram showing the distribution of distances of SMAD4 interacting regions in relation to the nearest transcriptional start site (TSS) at E9.5-E10.0 (25-30 somites) and E10.5 (34-38 somites). (B) The Average *Phastcons* conservation of the genomic regions enriched in SMAD4-chromatin complexes is shown at E9.5-E10.0 and E10.5. (C) Hierarchical clustering of the high-affinity matches for the top known and *de novo* motifs enriched in the SMAD4-bound regions at E9.5-E10.0 and E10.5. (D) The top five *de novo* motif identified in the genomic regions enriched in SMAD4-chromatin complexes in forelimb buds at E9.5-E10.0 and E10.5. (E) ChIP-qPCR validation of two previously known SMAD4 interacting genomic regions (Brugger et al., 2004; Korchynskyi and ten Dijke, 2002). Two biological replicates were analysed (mean ± standard deviation of three technical replicates). The left panels show the analysis for the BMP responsive element (BRE) regulating the *Id1* gene. The right panels show the analysis for the BRE regulating the *Msx2* gene.

### Identification of differentially expressed SMAD4 target genes

RNA-seq was used to identify the differentially expressed genes in the mesenchyme of wild-type and *Smad4*-deficient forelimb buds. As *Smad4*-deficient mouse embryos die prior to the onset of limb bud development, *Smad4* was conditionally inactivated in the forelimb bud mesenchyme using the *Prx1*-CRE transgene (*Smad4*^Δ/Δc^). *Prx1*-CRE-mediated *Smad4* inactivation results in clearance of mesenchymal SMAD4 proteins by around E10.0 (Bénazet et al., 2012). In contrast, SMAD4 proteins remain in the limb bud ectoderm, which circumvents the disruption of AER formation and massive mesenchymal apoptosis (Bénazet et al., 2009; Bénazet et al., 2012). As no mesenchymal apoptosis nor other gross-morphological abnormalities are observed in *Smad4*^Δ/Δc^ forelimb buds at the relevant forelimb bud stages comparative RNA-seq analysis was performed (Fig. 2, 3). First, pairs of age-matched wild-type and *Smad4*^Δ/Δc^ forelimb buds were analysed at E10.0 (30 somites) as by this developmental stage SMAD4 proteins have been cleared from the mutant mesenchyme and the SHH/GREM1/AER-FGF feedback signaling system is being established (Fig. 2A and Tables S1 and S2; Bénazet et al., 2012). Comparison of wild-type and *Smad4*^Δ/Δc^ forelimb buds identified a total of 668 differentially expressed genes (Fig.2B; DEGs: fold change ≥ .2; FDR <0.1). The cut-off was set at ≥ 2 to allow detection of spatial differences by whole mount RNA *in situ* hybridization (Probst et al., 2011). Among the 668 DEGs in early *Smad4*^Δ/Δc^ forelimb buds, 360 are up- and 308 down-regulated (Tables S1 and S2).

**Figure 2.**
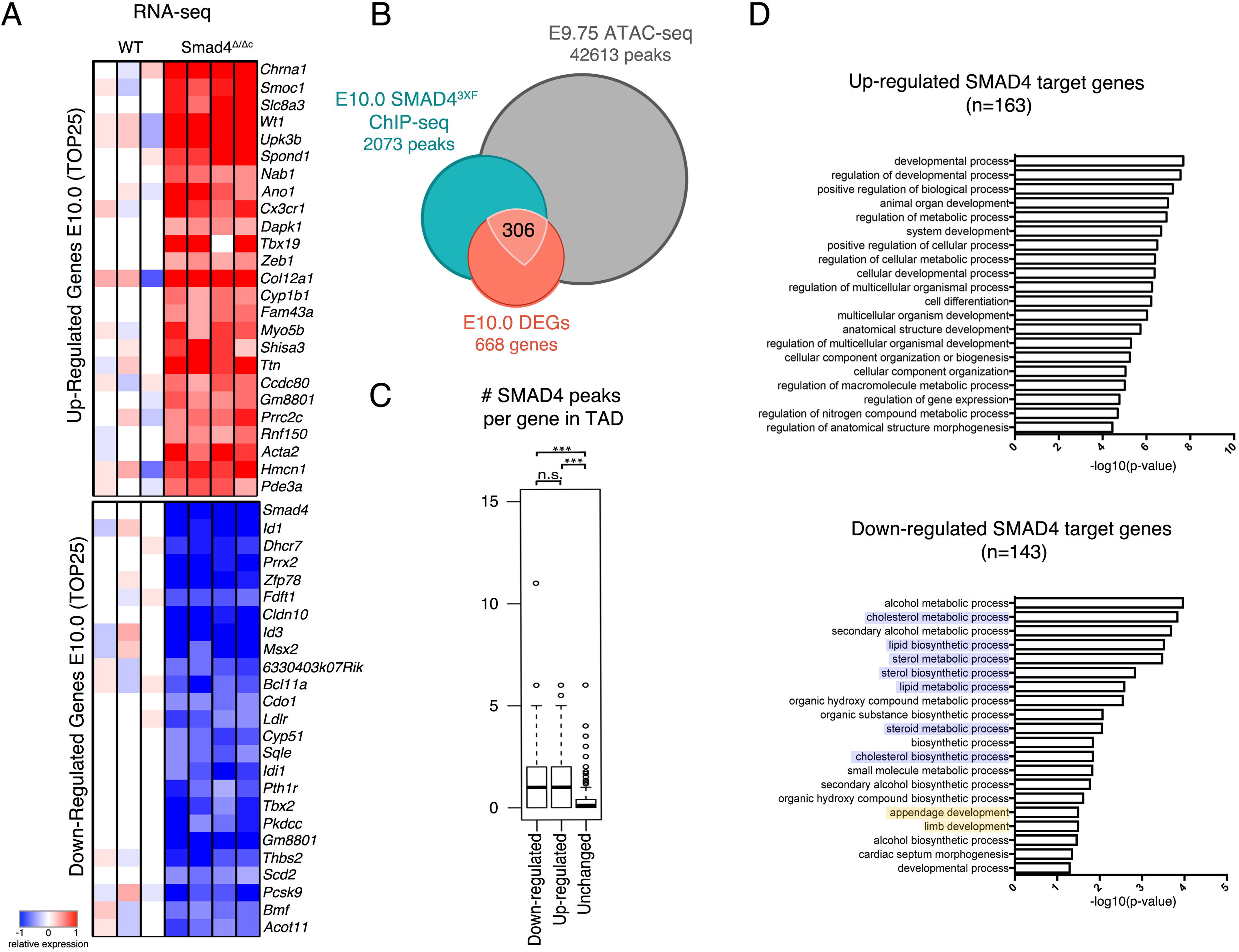
Comparative transcriptome analysis of wild-type and *Smad4*^Δ/Δc^ forelimb buds at E10.0 and identification of SMAD4 direct transcriptional targets. (A) Top 25 up- and down-regulated genes in *Smad4*^Δ/Δc^ forelimb buds at E10.0 (30 somites) are shown as heat maps. For each gene, the log2-ratio between the expression level in each sample and the mean (white) of the three biological replicates for the wild-type (WT) forelimb buds is shown. Red: increased expression, blue: reduced expression in comparison to the mean of the wild-type samples. Four biological replicates of *Smad4*^Δ/Δc^ forelimb buds were analysed. (B) Three-way Venn diagram showing the intersection between the significantly enriched ChIP-seq peaks (E9.5-E10.0, 25-30 somites), ATAC-seq peaks of age-matched wildtype forelimb buds (revealing the open chromatin regions, E9.75; 26 somites) and differentially expressed genes (DEGs) identified by RNA-seq analysis (E10.0, 30 somites). This intersection identifies 306 candidate SMAD4 target genes in limb buds at E9.5-E10.0. (C) Box plot representing the number of E9.5-E10.0 SMAD4^3xF^ ChIP-seq peaks within a topologically associated domain (TAD) of genes that are either up- or down-regulated in *Smad4*^Δ/Δ^ forelimb buds in comparison to genes with unaltered expression. Three asterisks: p≤1e-5 (Mann-Whitney test). (D-E) GO analysis of the up- and down-regulated SMAD4 target genes in *Smad4*^Δ/Δc^ forelimb buds. These target genes were defined by associating the genomic regions enriched in SMAD4-chromatin complexes with the closest up- or down-regulated gene in the *Smad4*^Δ/Δc^ transcriptome using a maximally 1Mb interval. Blue highlights GO terms for processes relevant to sterol biosynthesis and yellow highlights processes relevant to limb bud development (bottom panel).

**Figure 3.**
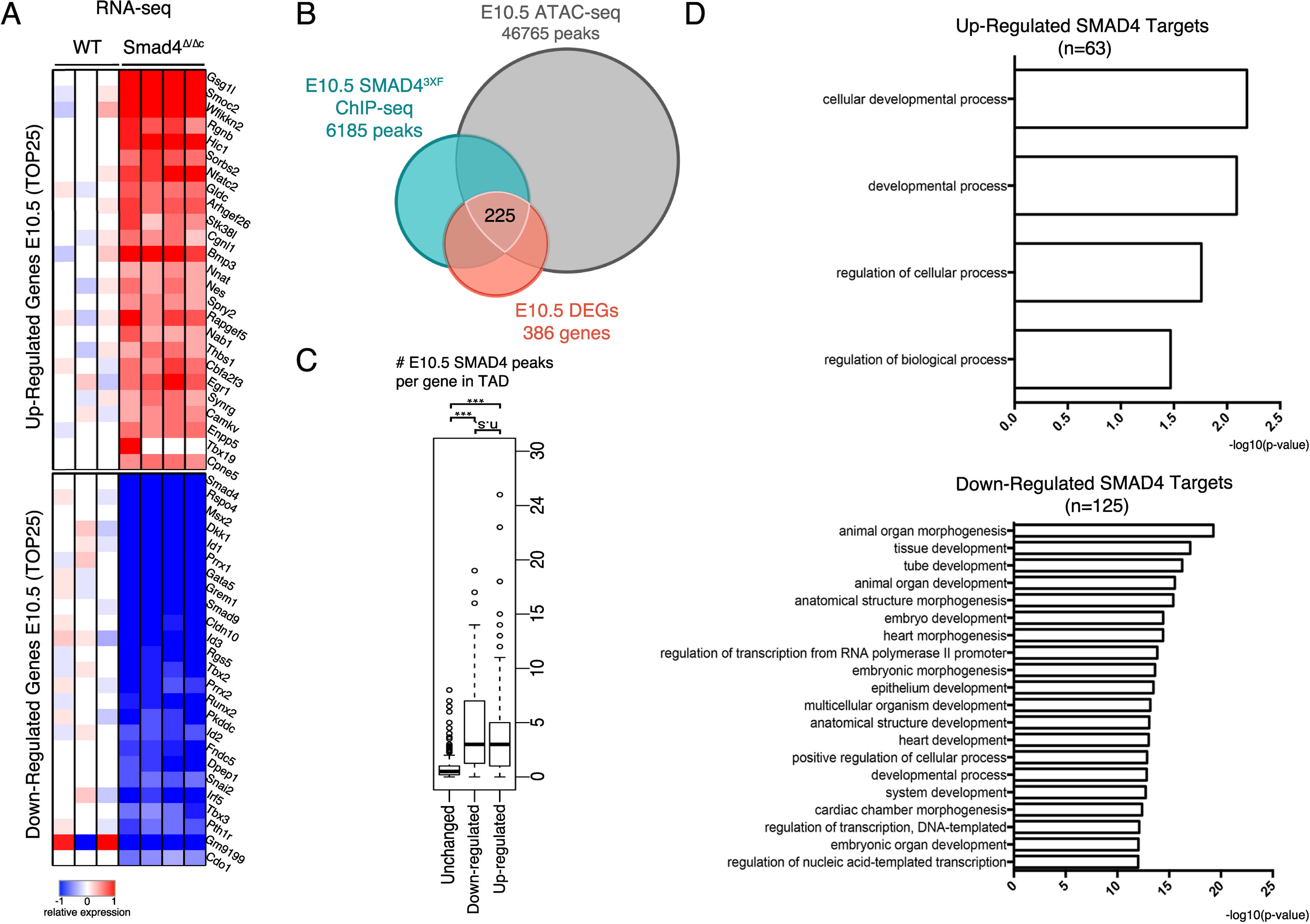
The range of SMAD4 target genes in forelimb buds at E10.5. (A) Top 25 down-regulated and up-regulated genes in *Smad4*^Δ/Δc^ E10.5 forelimbs (normalized to the mean expression in WT samples). The scale represents the logarithm of the ratio between the *Smad4*^Δ/Δc^ and mean WT expression levels. (B) Schematic representation of the intersection between the ChIP-seq (E10.5, 34-38 somites), ATAC-seq (E10.5, 35 somites) and RNA-seq (E10.5, 35 somites) datasets. The total numbers of ChIP peaks, ATAC peaks and DEGs are indicated. This intersection identifies 225 candidate SMAD4 target genes in forelimb buds at E10.5. (C) Box plot analysis representing the number of SMAD4^3xF^ ChIP-seq peaks within a topologically associated domain (TAD) harboring the SMAD4 target gene. Three asterisks: p≤1e5 (Mann-Whitney test) (D) GO enrichment analysis of biological processes for down- and up-regulated SMAD4 target genes.

The SMAD4 transcriptional target genes in early mouse forelimb buds (Fig. 2B) were identified as follows: the SMAD4 ChIP-seq peaks (E9.5-E10.0, 25-30 somites) overlapping regions of open chromatin as determined by ATAC-seq analysis in wild-type forelimb buds at E9.75 (26 somites, n=2) were assigned to the nearest DEG (E10.0, 30 somites) that is located within maximally a 1Mb genomic interval. This bioinformatics analysis identified 306 SMAD4 target genes that are differentially expressed in the mesenchyme of early mouse forelimb buds. Genes that are either up- or down-regulated in *Smad4*^Δ/Δc^ forelimb buds contain in general more SMAD4-binding regions within their topologically associating domains (TADs, Dixon et al., 2012) than genes whose expression is not changed (Fig. 2C). In *Smad4*^Δ/Δc^ forelimb buds at E10.0, the expression of 163 SMAD4 target genes is up-regulated (Table S3), while the others are down-regulated (n=143, Table S4). Gene Ontology (GO) analysis shows that genes with increased expression, i.e. target genes negatively regulated by SMAD4, function in various developmental processes, in agreement with the broad *Smad4* requirement during embryonic development (Figure 2D; Chu et al., 2004). GO analysis of SMAD4 target genes with reduced expression in mutant forelimb buds reveals an unexpected role of SMAD4 in the positive regulation of sterol/cholesterol biosynthesis and metabolism (blue shaded terms, Fig. 2E). As expected, this positively regulated set of SMAD4 target genes also includes genes that function during limb development (yellow shaded terms, Fig. 2E).

As BMP activity is progressively reduced due to increasing GREM1-mediated BMP antagonism (Bénazet et al., 2009), we used the same strategy to identify DEGs and SMAD4 target genes in forelimb buds at E10.5 (35 somites, Fig. 3 and Tables S5-S8). Comparison of wild-type and *Smad4*^Δ/Δc^ forelimb buds identified a total of 386 DEGs and 225 differentially expressed SMAD4 target genes (Fig. 3A, B). As for the earlier limb bud stage, more SMAD4-binding regions were detected in TADs of differentially expressed target genes than in the ones with no changes in expression in *Smad4*^Δ/Δc^ limb buds (Fig. 3C). This indicates that target genes whose limb bud expression depends critically on SMAD4 are regulated by the interaction of SMAD4-chromatin complexes with multiple rather than single *cis*-regulatory modules (CRMs, Fig. 3C). GO analysis of the SMAD4 target genes in forelimb buds at E10.5 points to functions in various developmental processes (Fig. 3D), but terms relevant to sterol/cholesterol biosynthesis are no longer enriched (compare to Fig. 2E). This indicates that SMAD4 up-regulates the expression of enzymes involved in cholesterol synthesis during the onset rather than during the progression of forelimb bud development.

### The limb bud mesenchymal *Smad4* deficiency causes premature transcriptional downregulation of cholesterol synthesis enzyme and intra-cellular cholesterol

The transcript levels of cholesterol biosynthesis enzymes (reviewed in Luo et al., 2020) is higher in wild-type than *Smad4*^Δ/Δc^ forelimb buds at E10.0 (Fig. 4A, B, Tables S2, S6). Only by E10.5, expression levels are reduced to a similar extent in both genotypes, which reveals the SMAD4 requirement for up-regulating/maintaining the transcription of cholesterol biosynthesis enzymes during the onset of limb bud development (prior to E10.5, Fig. 4B). Several of these downregulated enzymes are direct transcriptional targets of SMAD4 at E10.0 (7 of 16, Fig. 4A, B and Table S4). The target genes that are prematurely down-regulated in *Smad4*^Δ/Δc^ forelimb buds also include *Insig1, Ldlr*, *Pcsk9* and *Srebf1,* which are non-enzymatic regulators of the cholesterol pathway (Fig. 4B; Luo et al., 2020). Comparative WISH analysis shows that most cholesterol biosynthesis enzymes and regulators are expressed rather uniformly, which precludes detection of distinct spatial differences (Fig. S2). However, the spatial transcript distribution of key enzymes such as *Mvk, Idi1*, *Cyp51* and *Dhcr7* is altered in *Smad4*^Δ/Δc^ forelimb buds (Fig. 4C). Together with reduced transcript levels for most enzymes (left panels, Fig. 4B), this points to possible alterations in endogenous cholesterol biosynthesis in mutant limb buds at early stages (E10.0). The total cholesterol content includes both cell membrane-associated and intracellular cholesterol and a potential deficiency in endogenous cholesterol could be masked by exogenous cholesterol produced by other embryonic tissues or maternal origin (Tint et al., 2006). Therefore, the levels of intracellular cholesterol were analysed as follows: wild-type and *Smad4*^Δ/Δc^ limb bud mesenchymal progenitors (LMPs) were isolated from pairs of forelimb buds (E10.0; 28-30 somites) and cultured in cholesterol-free medium for 20-24 hours. After depletion of membrane-associated cholesterol, intracellular cholesterol levels were quantitated for LMPs of both genotypes (Table S9, for details see Materials and Methods and Vienken et al., 2017; Wilhelm et al., 2017). The primary values were used to determine the intracellular cholesterol levels per LMP (Fig. 4D). While there is inherent variability between LMPs from different forelimb buds, wild-type LMPs contain in average ∼4.6×10^−7^μg intracellular cholesterol, while these levels are reduced to ∼1.48×10^−7^μg cholesterol per cell in *Smad4*^Δ/Δc^ limb buds (Fig. 4D). This intracellular cholesterol deficiency is a likely consequence of the premature down-regulation of cholesterol biosynthesis enzymes in mutant forelimb buds (left panels, Fig. 4B). As cholesterol modification is required for SHH signalling and signal transduction (Li et al., 2006), we investigated potential alterations in cultured *Smad4*^Δ/Δc^ LMPs. However, cellular and biochemical analysis failed to reveal significant alterations, which is likely due to endogenous cholesterol synthesis being prematurely reduced by the *Smad4* deficiency rather than disrupted by e.g. inactivating the DHCR7 or the *Sc5d* enzymes (Fig. 4A; Cooper et al., 2003; Krakowiak et al., 2003). In agreement, no significant spatial changes in *Shh* expression and its targets *Gli1* and *Ptch1*, which serve as transcriptional sensors of SHH signal transduction are detected in *Smad4*^Δ/Δc^ forelimb buds (Fig. S3). In contrast, the SMAD 4 target gene *Hhip*, which encodes an inhibitor of SHH signaling (Chuang and McMahon, 1999), is upregulated in *Smad4*^Δ/Δc^ forelimb buds at E10.5 (Fig. S3, Table S5, S7).

**Figure 4.**
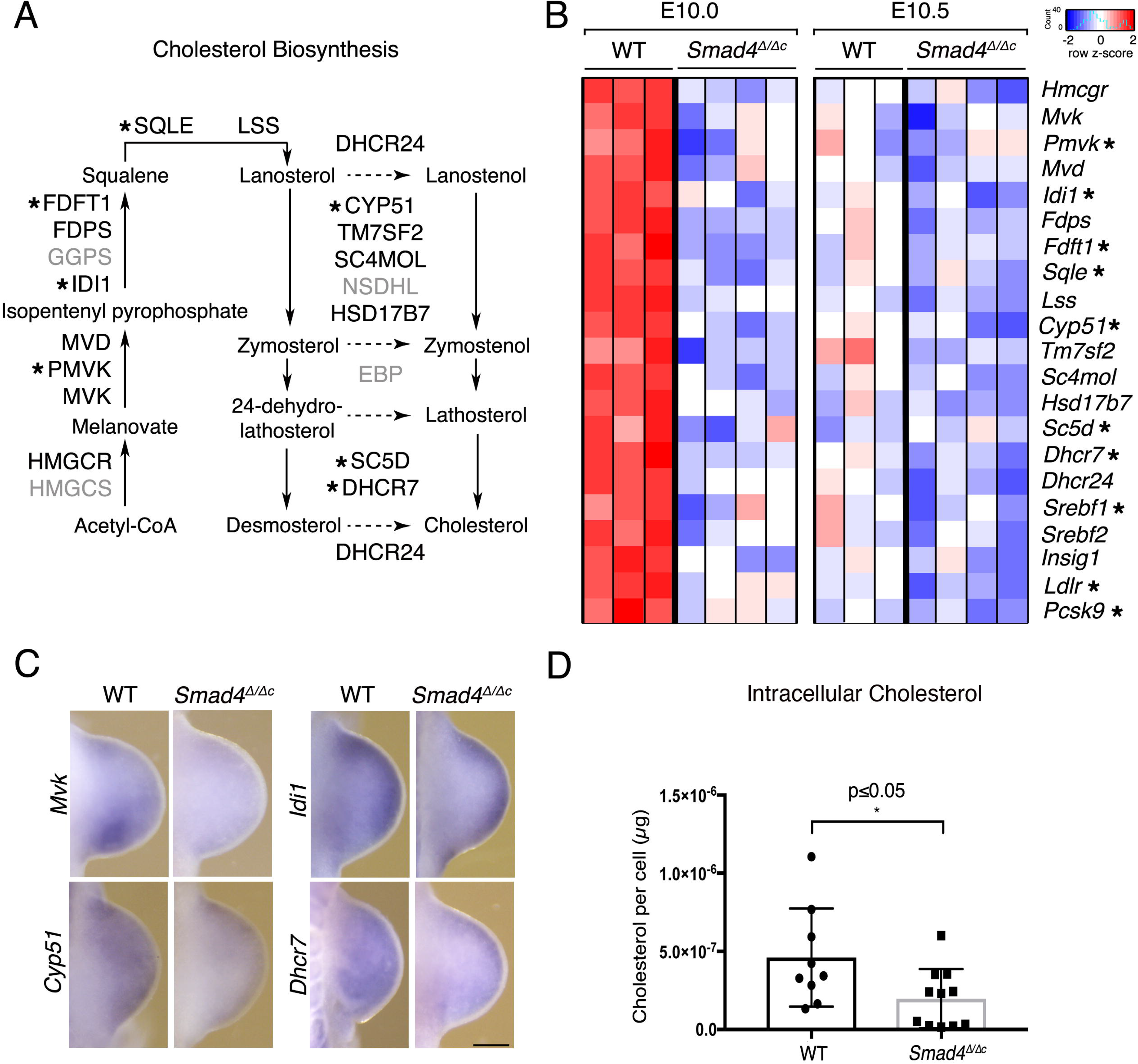
The expression of cholesterol biosynthesis enzymes is downregulated and intracellular cholesterol levels are reduced in *Smad4*^Δ/Δc^ limb buds. (A) Schematic representation of the cholesterol biosynthesis pathway. Enzymes down-regulated in *Smad4*^Δ/Δc^ forelimb buds at E10.0 are indicated in black, direct SMAD4 target genes are indicated by asterisks. Enzymes with unchanged expression are indicated in grey. (B) Heatmap showing expression of the downregulated genes encoding enzymes and additional regulators of cholesterol biosynthesis in wild-type and *Smad4*^Δ/Δc^ forelimb buds at E10.0 (30 somites) and E10.5 (35 somites). The heatmap indicates the Z-score of the read count per gene. Red indicates higher, blue lower than average expression levels. SMAD4 target genes are indicated by asterisks. (C) WISH analysis of key genes in the cholesterol biosynthesis pathway, whose spatial expression is clearly altered in *Smad4*^Δ/Δc^ forelimb buds at E10.0 (28-31 somites). Per gene and genotype n≥3 independent samples were analysed. Scale bar: 250µm. (D) Biochemical quantitation of intracellular cholesterol in wild-type (n=9) and *Smad4*^Δ/Δc^ LMPs (n=11 biological replicates) after culture in cholesterol-free medium (20-24 hrs). The intracellular cholesterol per cell was calculated by dividing the total cholesterol measured by the cell numbers for each sample (see Table S9). Intracellular cholesterol levels are lower in mutant than wild-type LMPs (p<0.05, Mann-Whitney test). The graphs show the individual data points plus the mean ± SD.

### The *Smad4* deficiency disrupts BMP signal transduction during early forelimb bud development

To gain insight into major molecular differences between wild-type and *Smad4*-deficient limb buds, the stage-specific and shared DEGs and SMAD4 target genes were identified (Fig. 5A, B). Not only is the number of DEGs and SMAD4 target genes reduced in forelimb buds at E10.5, but also few DEGs and SMAD4 target genes are shared between the two stages (E10.0: n=151; E10.5: n=43, Fig. 5A, B, Table S10-S13). Rather, most of the DEGs and SMAD4 target genes are markedly different during the onset (E9.5-E10.0) and progression of forelimb bud development (E10.5; Fig. 2, Fig. 3 and Fig. 5A, B). Interestingly, this change in the SMAD4 target genes correlates well with the observed differences in the enriched motifs in SMAD4-chromatin complexes (Fig. 1D) and parallels the shift from high to low BMP activity (Bénazet et al., 2009). As SMAD4 participates in both BMP and TGFβ signal transduction, we assessed the extent to which the expression of DEGs and inferred SMAD4 target genes (Figs. 2, 3) in the TGFβ and BMP pathways are altered between wild-type and *Smad4*^Δ/Δc^ forelimb buds (TGFβ GO:0007179 and BMP GO:0030509; Fig. 5C-5E). This analysis shows that a smaller fraction of genes assigned to the TGFβ than BMP pathway are differentially expressed (Fig. 5C). Only six of the 17 DEGs belonging to the TGFβ pathway are down-regulated in mutant forelimb buds, while others are precociously up-regulated (Fig. 5C, 5D). In contrast, analysis of the BMP pathway shows that the fraction of down-regulated DEGs increases during early limb bud outgrowth (from 11 to 15 of the 23 DEGs, Fig. 5C, 5E). This is intriguing as it parallels the reduction in BMP activity during progression of wild-type limb bud outgrowth, which does not occur in *Smad4*^Δ/Δc^ forelimb buds (Figure 5E; Bénazet et al., 2009; Bénazet et al., 2012). Furthermore the *Bmp2*, *-4*, *-7* ligands, which are required in the limb bud mesenchyme are upregulated, while the expression of transcriptional sensors for BMP signal transduction, *Msx2* and *Id1* is much reduced in *Smad4*^Δ/Δc^ forelimb buds (Fig. 5E, Fig. S4; Brugger et al., 2004; Lopez-Rovira et al., 2002). As no corresponding changes are detected in the *Tgf*β pathway (Fig. 5D, Fig. S4), *Smad4* functions predominantly or exclusively in BMP signal transduction during early limb bud development (E10.0-E10.5). The opposing effects of mesenchymal *Smad4*-deficiency on the expression of *Bmp* ligands and transcriptional sensors shows that BMP signal transduction is disrupted in the *Smad4* mutant mesenchyme. This is corroborated by the failure to upregulate mesenchymal *Grem1* via the feedback signaling system in response to increased *Bmp4* expression (Fig. 5E; Bénazet et al., 2009).

**Figure 5.**
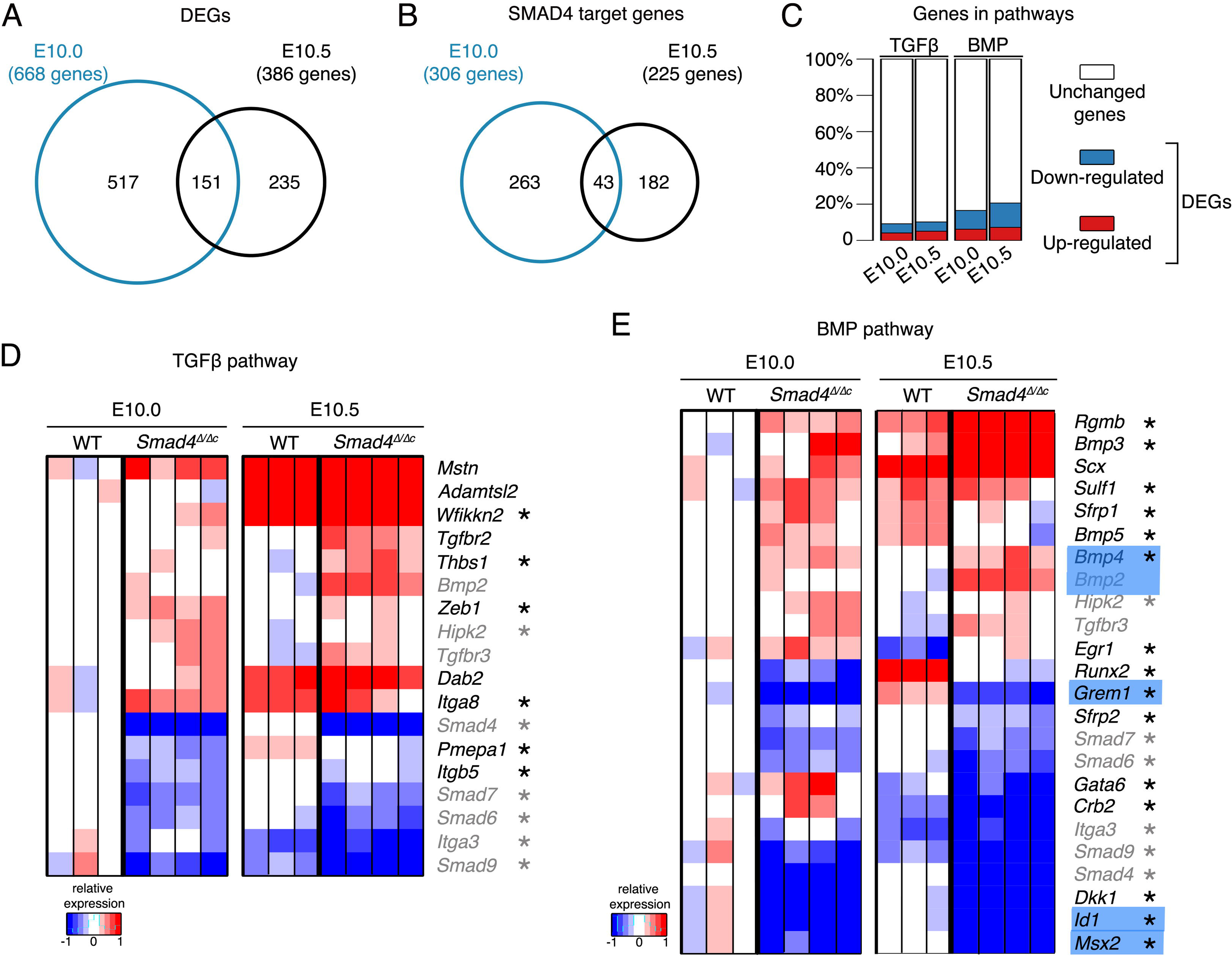
Differential regulation of gene expression in forelimb buds at E10.0 and E10.5, but SMAD4 predominantly impacts the BMP pathway at both stages. (A-B) Venn diagrams showing the intersection between the DEGs at E10.0 and E10.5 (A) and SMAD4 target genes at E10.0 and E10.5 (B). (C) Stacked bar plots show the percentage of DEGs functionally associated to the TGFβ (GO:0007179) and BMP (GO:0030509) pathways, respectively. (D-E) Heat maps showing the DEGs in the TGFβ (D) and BMP (E) pathways. For each gene, the log2-ratio between the expression level in each sample and the mean (white) of the three biological replicates for the wild-type (WT) forelimb buds are shown. Red: increased expression; blue: reduced expression in comparison to the mean of the wild-type samples. Genes indicated in black are either TGFβ or BMP pathway-specific, while genes indicated in grey are shared between the two pathways. The names of some key genes in the BMP pathway are underlaid in blue - for details see results. Asterisks mark the SMAD4 target genes among the DEGs.

### *Smad4* controls the spatial up-regulation of target genes by interacting with CRM enhancers active in the anterior forelimb bud mesenchyme

Whole mount RNA *in situ* hybridization (WISH) was used to analyse the spatial distribution of SMAD4 target genes, whose expression is reduced in *Smad4*^Δ/Δc^ forelimb buds (E10.0, Fig. 2A; Table S4). Of 143 down-regulated SMAD4 target genes, 91 could be analysed by comparative WISH of wild-type and mutant forelimb buds at E10.0 (29-31 somites) and the genes analysed further are shown in Fig. 4C and Fig. 6 (see also Fig. S2, Fig. S3). This screen revealed the reduced expression (Fig. 6A) and spatial alterations in the anterior mesenchyme of *Smad4*^Δ/Δc^ forelimb buds for target genes functioning in the BMP pathway (Fig. 6B, C). They include several regulators of BMP signalling in limb buds such as the transcriptional sensors *Id1, Id2* and *Id3*, *Msx2* and the inhibitory SMAD proteins *Smad6* and *Smad7* proteins (Figure 6B; Zhao et al., 2000). *LacZ* reporter assays establish that for three of these BMP pathway genes (*Id1*, *Id2* and *Msx2*), the genomic regions enriched in SMAD4-chromatin complexes encode *bona*-*fide* CRMs that function as transcriptional enhancers (Fig. 6C). The spatial activities of these enhancers recapitulate significant aspects of limb bud mesenchymal expression of the associated *Id1*, *Id2* and *Msx2* target genes (Fig. 6B, C). The other SMAD4 target genes, whose spatial expression is reduced and altered in *Smad4*^Δ/Δc^ forelimb buds at early stages (Fig. 6A, D) function either in antero-posterior limb bud patterning (*Alx4*, *Tbx2*; Farin et al., 2013; Kuijper et al., 2005), outgrowth and/or chondrogenesis (*Sfrp2*, *Snai1*, *Lhx2* and *Prrx2*; Chen and Gridley, 2013; Geetha-Loganathan et al., 2008; Taher et al., 2011; Tzchori et al., 2009). With the exception of *Mxd4* and *Sfrp2*, these genes are part of the differentially expressed target genes shared between *Smad4*^Δ/Δc^ forelimb buds at E10.0 and E10.5 (n=42, Fig. S5). In particular, the early and persistent downregulation of SMAD4 target genes functioning in the BMP pathway (*Smad6*,-*7*, *Id1*,-*2*,-*3*, *Msx2*) and bone development (*Lhx2, Snai1, Pkdcc, Pthr1*; Imuta et al., 2009; Karaplis et al., 1994; Probst et al., 2013) indicates that the disruption of chondrogenesis and bone formation (Bénazet et al., 2012) is rooted in these early transcriptional changes (Fig. 6D, Fig. S5). The SMAD4-enriched CRMs associated with the *Alx4*, *Lhx2* and *Pkdcc* genes display robust and predominant enhancer activities in the anterior limb bud mesenchyme (bottom panels, Fig. 6D). These enhancer activities together with the transcriptional downregulation of the associated SMAD4 target genes in the anterior mesenchyme of *Smad4*^Δ/Δc^ forelimb buds (arrow heads in Fig. 6B, 6D) show that one of the essential SMAD4 functions is the upregulation of target genes through enhancers active in the anterior mesenchyme (Fig. 6C, bottom panels Fig. 6D).

**Figure 6.**
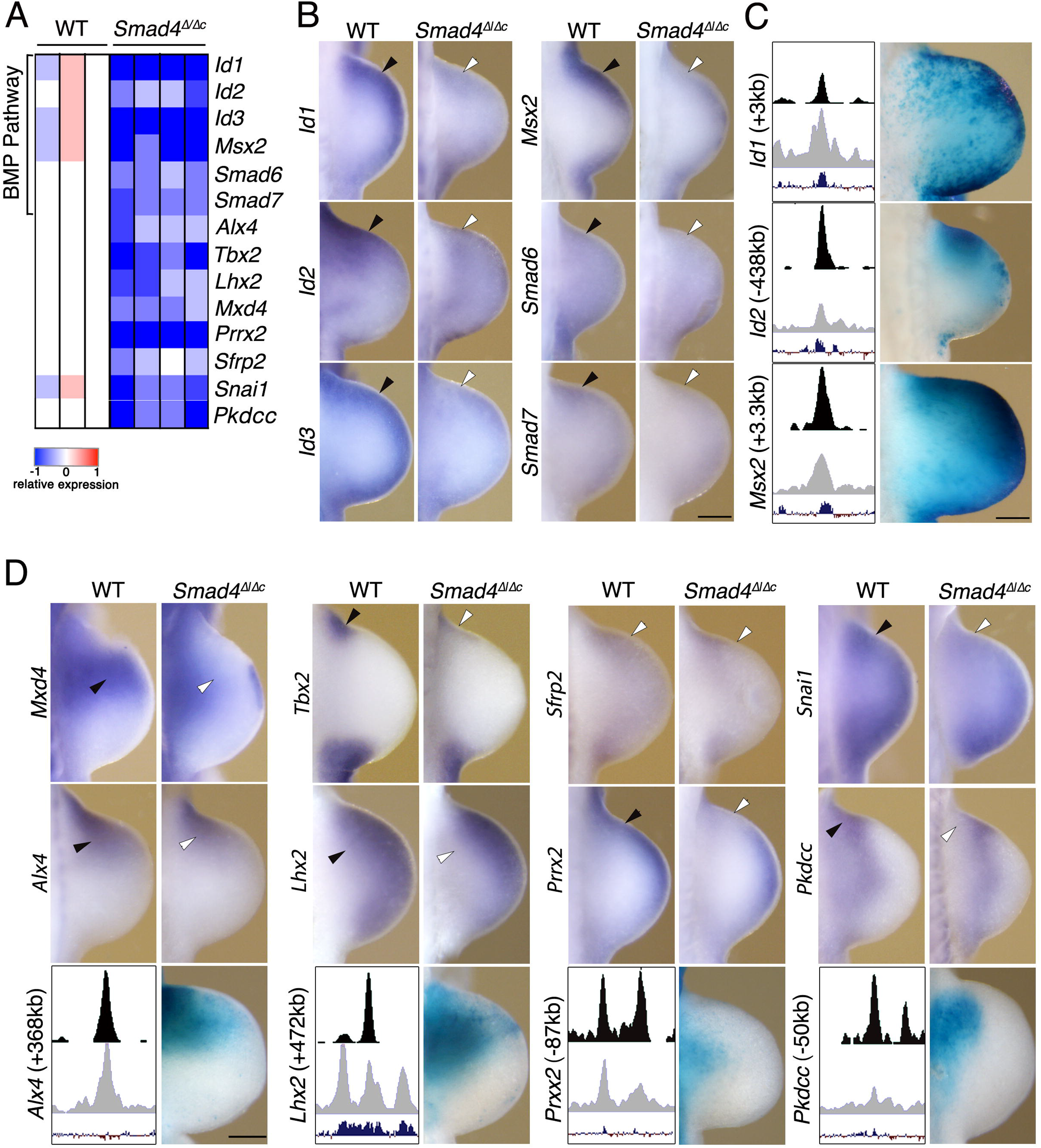
Target genes positively regulated by SMAD4 in the anterior forelimb bud mesenchyme. (A) Heat map of the target genes, whose expression is positively regulated by SMAD4 in the anterior forelimb bud mesenchyme. For each gene, the log2-ratio between the expression level in each sample and the mean (white) of the three biological replicates for wild-type (WT) forelimb buds is shown. Red: increased expression; blue: reduced expression in comparison to the mean of the wild-type samples. (B) Comparative WISH analysis of selected BMP pathway SMAD4 target genes in WT and *Smad4*^Δ/Δc^ forelimbs at E10.0 (28-31 somites). (C) Analysis of the *LacZ* reporter activity of SMAD4-enriched candidate CRMs associated with the *Id1*, *Id2* and *Msx2* target genes. Left panel shows a scheme depicting the genomic region harbouring the CRM with the SMAD4 ChIP-seq peak (top), the ATAC-seq peak (open chromatin, middle) and the evolutionary conservation (bottom). The right panel shows the expression of the CRM-*LacZ* reporter in forelimb buds. (D) SMAD4 target genes, whose spatial expression in the anterior limb bud mesenchyme is altered in *Smad4*^Δ/Δc^ forelimbs at E10.0. Right-hand lower panels show the activity of the SMAD4-enriched candidate CRMs in the *Alx4* and *Lhx2* loci in the anterior forelimb bud mesenchyme at E10.5. Panels (B-D): WISH: n≥3 independent samples were analysed per gene and genotype. For each CRM-*LacZ* reporter construct n≥3 independent founder embryos were obtained and the most representative patterns are shown. Scale bars: 250µm. (E) The anterior SMAD4 target gene network: interactions between SMAD4 target genes were identified using Ingenuity Pathway Analysis (grey arrows) in combination with manual curation (black arrows).

### SMAD4-controlled gene regulatory networks co-regulated by SHH signaling

This molecular analysis started to uncover the SMAD4-regulated GRNs (Fig. 3, 6) that function during the transient early patterning phase that specifies the digits in the forelimb bud mesenchyme (Zhu and Mackem, 2011; Zhu et al., 2008). During this early phase, SHH signalling is required to coordinate antero-posterior and proximo-distal limb bud patterning (AP and PD patterning; Probst et al., 2011; reviewed in Zuniga, 2015). Therefore, the extent to which SMAD4-regulated genes are co-regulated by SHH signalling was determined by comparative analysis of *Smad4* and *Shh* DEGs (Probst et al., 2011) during early mouse limb bud development (E10.0 to E10.5, Fig. 7A,B and Table S14). 111 shared DEGs were identified and among them 65 are SMAD4 target genes and 37 are required for normal limb development (Tissieres et al., 2020). These results indicate that SMAD4 and SHH signalling co-regulate GRNs with essential functions in early mouse limb buds. Strikingly, the majority of the shared DEGs are regulated in a discordant manner by the two pathways (79 genes; Fig. 7A; Table S14) and more than half are up-regulated in *Smad4* deficient limb buds (upregulated: 43, downregulated: 36, Fig. 7A; Table S14). Among the SMAD4 target genes 7 are BMP pathway genes regulated in a discordant manner, which includes the BMP antagonist *Smoc1* (Fig. 7A; see also Fig. 5E and Fig. S6; Okada et al., 2011; Thomas et al., 2017). Furthermore about half of all discordantly regulated genes are transcription factors, pointing to an important amplification of the response to BMP and SHH signalling in early limb buds. In contrast, few genes with known functions in early limb bud patterning and outgrowth are present among the 32 DEGs that are concordantly regulated by both pathways (Fig. 7B, Table S14). Notable exceptions are *Grem1* and some cholesterol pathway genes (*Dhcr7*, *Dhcr24*, *Insig1* and *Cyp1b1*), which are downregulated in both *Smad4* and *Shh*-deficient limb buds (Fig. 7B; Table S14; see also Fig.4).

**Figure 7.**
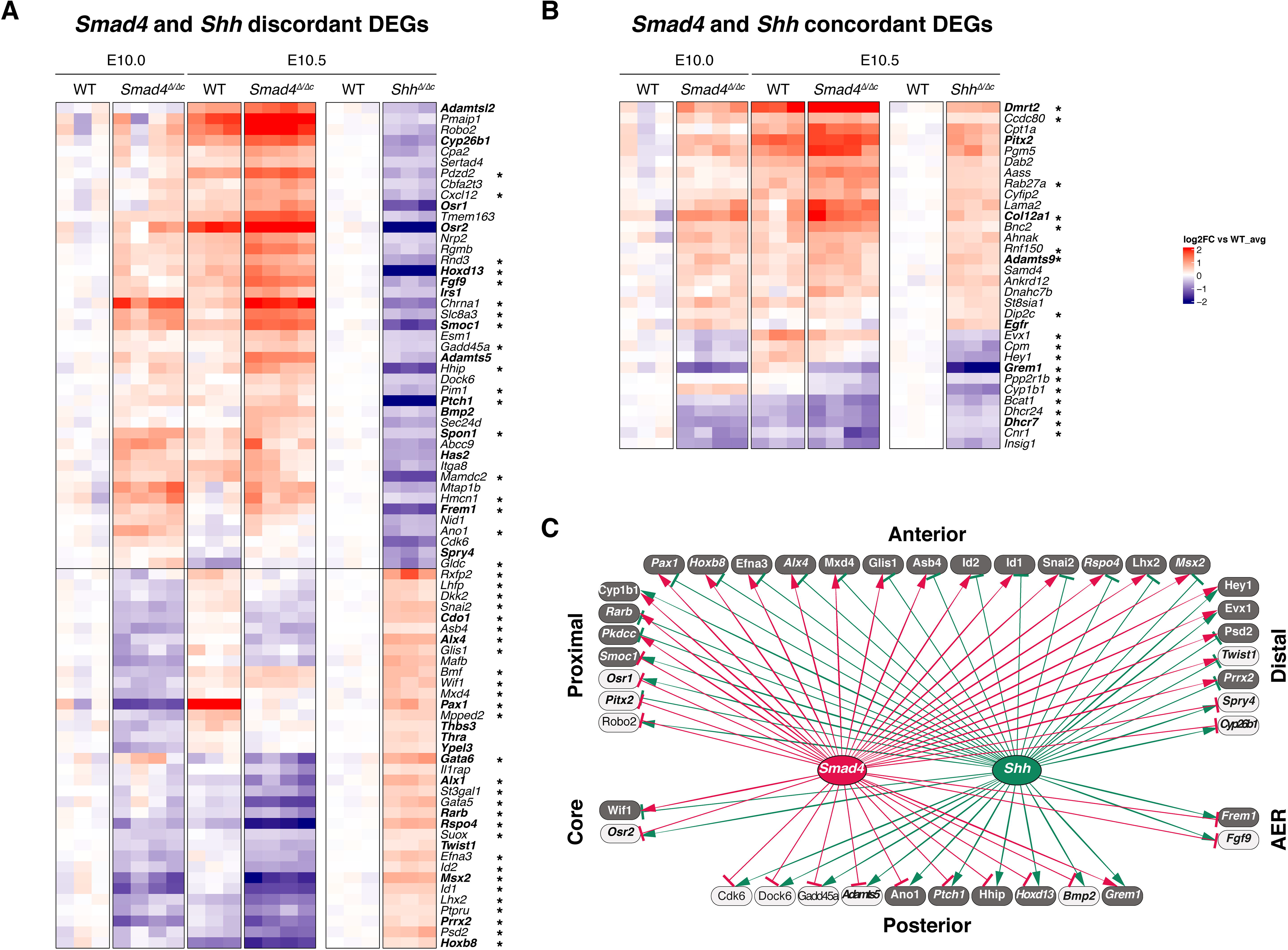
Antagonistic SMAD4 and SHH pathway interactions control antero-posterior patterning during progression of limb development. (A), (B) Heatmaps showing the differentially expressed genes (DEGs) identified by comparing WT and *Smad4*^Δ/Δc^ transcriptomes at E10.0 and E10.5 (WT: n=3, *Smad4*^Δ/Δc^: n=4 biological replicates) and WT and *Shh* ^/^ transcriptomes at E10.5 (WT: n=3, *Shh*^Δ/Δ^ : n=3 biological replicates) respectively. DEGs showing a fold change ≥1.2 and FDR<0.1 were analyzed (Table S14). Asterisks mark the SMAD4 target genes among the DEGs and genes in bold are associated to limb mutations. (A) *Smad4* and *Shh* discordant DEGs. (B) *Smad4* and *Shh* concordant DEGs. (C) The shared *Smad4* and *Shh* GRN consisting of genes with distinct spatial expression in mouse limb buds. The shared *Smad4* and *Shh* DEGs were grouped according to their spatial “anterior”, “posterior”, “proximal” or “distal” expression bias in the limb bud mesenchyme (see main text). In addition, two DEGs are expressed in the “core” mesenchyme without an apparent AP or PD bias and two in the AER (Table S15). The *Smad4* and *Shh* interactions within GRN are indicated by red and green lines, respectively. Positive interactions (DEGs downregulated in *Smad4* or *Shh*-deficient limb buds) are represented by arrows and negative interactions (DEGs upregulated) are represented by inhibitory arrows. DEGs shown as dark grey boxes are direct SMAD4 target genes. Genes indicated in italics have been linked to mutations causing limb skeletal phenotypes.

To gain insight into the interactions of these two pathways during early limb bud patterning, we screened the shared DEGs (Fig. 7A, B) for genes with distinct spatial expression patterns in mouse limb buds using the Mouse Genome Informatics (http://www.informatics.jax.org) and EMBRYS databases (https://www.embrys.jp). This analysis identified 41 genes with spatial restricted and asymmetrical distributions in early limb buds. According to their spatial distribution, DEGs were categorized as anteriorly, posteriorly, proximally or distally expressed genes. The more widely expressed genes were assigned to the category matching the limb bud mesenchymal region of their predominant/highest spatial expression (column “expression bias” in Table S15). For example *Msx2* is expressed anteriorly and posteriorly in wildtype mouse limb buds at E10.5, but as the anterior domain is wider, *Msx2* was annotated as an anteriorly biased gene (Fig. 6B, Table S15). In contrast, *Pkdcc* was annotated as a proximal gene because it is expressed throughout the proximal mesenchyme and excluded from the distal limb bud mesenchyme (Fig. 6D, Table S15). Strikingly, all anteriorly biased genes are SMAD4 target genes (dark grey boxes, Fig. 7C), whose expression is positively regulated by *Smad4* but negatively regulated by *Shh* (Fig. 7C; *Smad4*: red activating arrows and *Shh*: green inhibitory arrows). Conversely, the majority of posteriorly biased genes are negatively regulated by *Smad4* (red inhibitory arrows) and positively regulated by *Shh* (green activating arrows, Fig. 7C). The exception is *Grem1*, which is positively regulated by both SMAD4-mediated BMP signal transduction and SHH signaling as part of the self-regulatory feedback signaling system (Fig. 7C; Bénazet et al., 2009; Bénazet et al., 2012). In addition, *Smad4* and *Shh* have opposing effects on the majority of DEGs with asymmetrical distributions along the PD axis. In contrast to the AP limb bud axis, there appears to be no predominant positive or negative regulatory impact on PD gene expression from either pathway in early limb buds (Fig. 7C). This analysis indicates that SMAD4-mediated BMP and SHH signalling have overall rather opposing effects on co-regulated genes that are part of the GRNs regulating AP and PD axes patterning (reviewed in Zuniga and Zeller, 2020) during early mouse limb bud outgrowth.

## DISCUSSION

The present study identifies the SMAD4 target genes in the early mouse forelimb bud mesenchyme. SMAD4-mediated BMP signal transduction is predominant during the onset of forelimb bud development (Pignatti et al., 2014). In particular, the *Bmp2, -4,* - *7* ligands are expressed at high levels and essential for forelimb bud patterning and skeletal development (Bénazet et al., 2009; Luo et al., 1995). The *Tgf*β*2* and *Tgf*β*3* ligands are also expressed at comparable levels in early forelimb buds (this study; reviewed in Lorda-Diez et al., 2021), however inactivation of *Tgf*β*r2* showed that TGFβ signalling is not essential for limb bud patterning (Dunker et al., 2002). In contrast, limb bud mesenchymal *BmprI* inactivation cause patterning defects and SMAD4-mediated BMP-signal transduction is essential for initiation of chondrogenesis (Bénazet et al., 2012; Lim et al., 2015; Ovchinnikov et al., 2006). While crosstalk between BMP and TGFβ SMAD4-mediated signal transduction is possible (Karamboulas et al., 2010), SMAD4 is specifically required for BMP-signal transduction in early limb buds as the expression of *Bmp* ligands is up-regulated in *Smad4*-deficient limb buds, while the up-regulation of known targets and sensors of BMP activity such as *Id* genes, *Msx2* and *Grem1* are disrupted (this study; Bénazet et al., 2009; Bénazet et al., 2012; Brugger et al., 2004; Hsu et al., 1998; Lopez-Rovira et al., 2002). While BMP activity is high during initiation of limb bud development, it drops to lower levels during outgrowth and rapid proliferative expansion of LMPs (Reinhardt et al., 2019). This reduction is paralleled by significant changes in the binding specificities of SMAD4-chromatin complexes and range of DEGs and SMAD4 target genes in the forelimb bud mesenchyme (this study). However, SMAD4 target genes expressed in the anterior forelimb bud mesenchyme are likely targets of BMP4 and/or BMP7 signal transduction as their expression overlaps these genes in the anterior mesenchyme.

Unexpectedly, SMAD4 is required for maintaining the expression of the majority of cholesterol biosynthesis enzymes in early limb buds as their expression is prematurely down regulated and intracellular cholesterol reduced in *Smad4*^Δ/Δc^ forelimb buds. In addition to maternal sterols being a major source of cholesterol, defects in embryonic cholesterol synthesis cause congenital malformations similar to *Shh* loss-of-function defects (reviewed in Porter and Herman, 2011). It has been shown that the active SHH signaling peptide and SMO-mediated signal transduction depend on cholesterol modification, whose synthesis requires twenty different enzymes (reviewed in Radhakrishnan et al., 2020). Extensive follow-up analysis did neither reveal reproducible and significant alterations in the response to *Shh* signaling in *Smad4*^Δ/Δc^ forelimb buds, nor clearly identify cholesterol-dependent alterations in SHH and SMO-mediated signal transduction in *Smad4*^Δ/Δc^-deficient LMPs. This is likely due to the fact that the *Smad4* deficiency reduces, but does not disrupt endogenous cholesterol biosynthesis in contrast to mutations in the *DHCR7* enzyme (reviewed in Horvat et al., 2011). The reduction in endogenous cholesterol synthesis is likely compensated by maternal cholesterol (Tint et al., 2006) or cholesterol from other embryonic tissues as the *Smad4* inactivation is limb bud mesenchyme specific (Bénazet et al., 2012 and this study). However, our analysis shows that SMAD4 negatively regulates the expression of the SHH receptor *Ptch1* and the extracellular SHH antagonist *Hhip*, which might contribute to modulating the range of SHH signal transduction in limb buds (reviewed in Briscoe and Therond, 2013). In particular, increased *Hhip* expression might reduce SHH activity in the more central and anterior mesenchyme of *Smad4*^Δ/Δc^ forelimb buds (Chuang et al., 2003 and this study). The transcriptional regulation of cholesterol enzymes may be a more general function of SMAD4-mediated signalling as during palate development, transcriptional profiling identified a down-regulation of cholesterol synthesis enzymes and an upregulation of *Hhip* in *Tgfbr2-*deficient mouse embryos (Pelikan et al., 2013). BMP signalling functions in multiple processes during temporal progression of limb bud development, which includes AER formation, limb bud outgrowth, chondrogenesis and ultimately induction of mesenchymal apoptosis to separate the digits (Pignatti et al., 2014). Based on loss-of-function genetic and gain of function studies, it was proposed that the BMP pathway functions downstream of SHH signaling in the limb bud mesenchyme, but at the same time inhibits SHH signalling during limb bud patterning and outgrowth (Bastida et al., 2009; Bénazet et al., 2009; Bénazet et al., 2012; Panman et al., 2006; Selever et al., 2004; Zuniga et al., 1999). Inactivation of *Bmp* ligands and their receptors in the limb bud mesenchyme causes both pre- and postaxial polydactylies in agreement with *Bmp* expression in the anterior and posterior distal mesenchyme (Bandyopadhyay et al., 2006; Bénazet et al., 2009; Katagiri et al., 1998; Selever et al., 2004). However, the resulting limb skeletal phenotypes could be far downstream or indirect effects of disrupting BMP-signal transduction as a consequence of global alterations affecting morphogenetic signaling during limb bud development. Therefore, the identification of SMAD4 targets genes with morphoregulatory functions in early limb buds provides important insights into the direct impact of SMAD4 on gene expression (this study). SMAD4 regulates the expression of genes in both the anterior (including *Alx4*, *Tbx2*, *Msx2, Prxx2, Snai1*, and posterior limb bud mesenchyme (including Hoxd13, Grem1, Bmp2, Cdk6; this study and Bénazet et al., 2012; Lopez-Rios et al., 2012). Together with previous studies, our analysis reveals the direct involvement of SMAD4-mediated BMP signal transduction in the overall transcriptional upregulation of anterior and downregulation of posterior genes during early limb bud patterning (this study and Bandyopadhyay et al., 2006; Bastida et al., 2009; Bénazet et al., 2009; Bénazet et al., 2012; Ovchinnikov et al., 2006; Selever et al., 2004). Furthermore, the SMAD4 target genes *Alx4, Msx2, Prrx2* and *Dkk1* are required to restrain the developing limb bud to pentadactyly as their genetic inactivation causes polydactyly (Berge et al., 1998; Kuijper et al., 2005; Lallemand et al., 2005; Mukhopadhyay et al., 2001). This indicates that SMAD4-mediated BMP signal transduction overlapping the transient early phase of SHH-mediated digit patterning (Zhu et al., 2008) is required to maintain pentadactyly by direct transcriptional regulation of its target genes, many of which are also regulated by SHH signaling. SHH is not only essential for AP digit patterning but also functions in coordinating AP and PD axes development during limb bud outgrowth (Probst et al., 2011; Zhu et al., 2008).This is of interest in light of the globally opposing transcriptional regulation by SMAD4 and SHH signaling, which reveals direct antagonistic interactions in regulating the transcription of genes with AP and PD expression bias and functions. SHH signaling by the polarising region up-regulates gene expression in the posterior mesenchyme but also prevents posterior expansion of anterior SMAD4 target genes. In the anterior mesenchyme, BMP signals via SMAD4 to upregulate and/or maintain the expression of SMAD4 target genes independently of SHH signaling, while preventing anterior expansion of posterior genes with the exception of the BMP antagonist *Grem1*. *Grem1* is a transcriptional node in the self-regulatory signaling system that integrates transacting inputs from both SMAD4-mediated BMP and GLI-mediated SHH signal transduction into transcriptional up-regulation and spatial regulation of *Grem1* expression (Malkmus et al., in press). The increase in GREM1-mediated BMP antagonism is balanced by feedback regulation, which results in persisting low levels of BMP activity within the posterior limb bud mesenchyme (Bénazet et al., 2009; Malkmus et al., in press). Low level BMP activity is relevant to restrain the autopod to pentadactyly as transgene-mediated *Grem1* overexpression results in polydactyly (Berge et al., 1998; Kuijper et al., 2005; Lallemand et al., 2005; Mukhopadhyay et al., 2001; Norrie et al., 2014). Most importantly, GREM1-mediated BMP antagonism in the posterior mesenchyme is required to establish and propagate the self-regulatory feedback signaling system that enables distal progression of limb bud outgrowth and patterning (Zuniga, 2015; Zuniga et al., 1999). The intricacy of the direct molecular interactions is further exemplified by the fact SMAD4 directly controls the transcription of two of the three BMP ligands, *Bmp2* and *Bmp4* and the BMP antagonists *Grem1* and *Smoc1* that are essential for normal limb bud development. The analysis of SMAD4-regulated genes (DEGS) and target genes leads us to propose that BMP activity is maintained at high levels in the anterior mesenchyme, which is supported by the lack of BMP antagonist expression in this territory (*Smoc1* and *Grem1*, this study and Okada et al., 2011; Zuniga et al., 1999). During pectoral fin bud development, BMP signalling gradients are important for fin bud growth and these gradients are modulated by SMOC1, which reinforces the importance of tight modulation of BMP activities during morphogenesis (Mateus et al., 2020).

Mouse genetic experiments and experimental manipulation of chicken limb buds have shown that tampering with BMP activity levels alters *Shh* expression (Bastida et al., 2009; Bénazet et al., 2009; Bénazet et al., 2012; Norrie et al., 2014). Our analysis shows that this is likely indirect as *Shh* expression and signal transduction are not affected in *Smad4*-deficient mouse limb buds. Rather unexpectedly, our study identifies the Hedgehog inhibitory protein *Hhip* as a negatively regulated SMAD4 target gene, which indicates that SMAD4-mediated signal transduction not only modulates BMP activity but also participates in finetuning the SHH activity range in limb buds. Thus SMAD4-mediated BMP signal transduction and SHH signaling have both direct opposing impacts (posterior mesenchyme) and coordinated interactions (anterior limb bud mesenchyme; Bastida et al., 2009; (Bénazet et al., 2009; Dunker et al., 2002). This highlights the intricate control of morphogenetic signaling by BMPs and SHH during the early phase critical to digit specification and patterning in mouse limb buds (Zhu et al., 2008). In the neural tube, high BMP activity is required for dorsal neural tube patterning while ventrally, SHH and the BMP antagonist chordin are co-expressed in the notochord, and SHH and the interaction of the two secreted factors is required to induce the ventral floor plate (Patten and Placzek, 2002). In fact, *Shh* and *Bmp* ligands are co-expressed in numerous tissues, which indicates that the interactions of the two pathways could be conserved during embryogenesis (Bitgood and McMahon, 1995). Functionally relevant BMP and SHH signaling interactions have been reported for other developing tissues such as tooth and mandibular arch development (Harris et al., 2002; Li et al., 2015; Madison et al., 2005; Xu et al., 2019). In contrast to these previous studies, we establish that a large fraction of the genes in the shared SMAD4-SHH GRN are direct transcriptional targets of SMAD4-mediated BMP signal transduction during the early phase of limb bud patterning and outgrowth.

## MATERIALS AND METHODS

### Ethics statement, mouse strains and embryos

All experiments conducted with mice and embryos of both sexes at the developmental ages indicated were performed strictly respecting Swiss laws, the 3R principles and the principles of the Basel Declaration. All animal studies were evaluated and approved by the Regional Commission on Animal Experimentation and the Cantonal Veterinary Office of the city of Basel (license 1950). To conditionally inactivate *Smad4* in the forelimb bud mesenchyme, the *Prx1-Cre* strain was used (Logan et al., 2002; Yang et al., 2002). *Prx1-Cre^Tg/Tg^Smad4*^Δ^*^/+^* males were crossed with *Smad4^flox/+^* females to obtain experimental embryos that carry a constitutive *Smad4* null allele and a conditionally inactivated *Smad4* allele (*Prx1- Cre^Tg/+^Smad4*^Δ/Δ^, referred to as *Smad4*^Δ/Δ^). Resulting *Prx1-Cre Smad4* littermates were used as controls for all experiments (referred to as “wild-type” in the text). *Shh*^Δ/Δ^ embryos were obtained from crossing mice heterozygous for the *ShhCre* allele (Harfe et al., 2004)

### Generation of the *Smad4*^3xF^ mouse allele

The *Smad4*^3xF^ mouse strain was generated by introducing a 3xFLAG epitope tag in the endogenous SMAD4 protein by conventional homologous recombination in mouse ES cells. The used targeting vector consisted of two homology arms flanking the 3’-end of *Smad4* coding sequence, in which the 3xFLAG was inserted in frame between the exon 12 and the 3’UTR. A floxed *neomycin* selection cassette was inserted downstream of *Smad4* locus in the 3’ homology arm. This targeting vector was linearized and electroporated in R1 embryonic stem (ES) cells to generate the *Smad4*^3xF-Neo^ allele. ES cells clones were screened for correct recombination by Southern Blot analysis using probes located either outside of the 5’ or 3’ homology arms (5’ and 3’probes) and for the *neomycin* cassette to exclude ES cell clones with random integration of the targeting construct. Correctly targeted ES cells clones were then injected in C57BL/6 blastocysts by the Centre of Transgenic Mice (CTM) of the University of Basel. Chimeric males were obtained from three independent ES cell clones and mated with *CMV-*Cre females (C57BL/6 background) to delete the floxed *neomycin* selection cassette. Germline transmission was assessed using PCR genotyping. *CMV*-Cre-mediated deletion of the floxed *neomycin* selection cassette generated the *Smad4^3xF^* allele. Specific primers were used to discriminate between the *Smad4^+/+^, Smad4^3xF-Neo^* and *Smad4^3xF^* alleles.

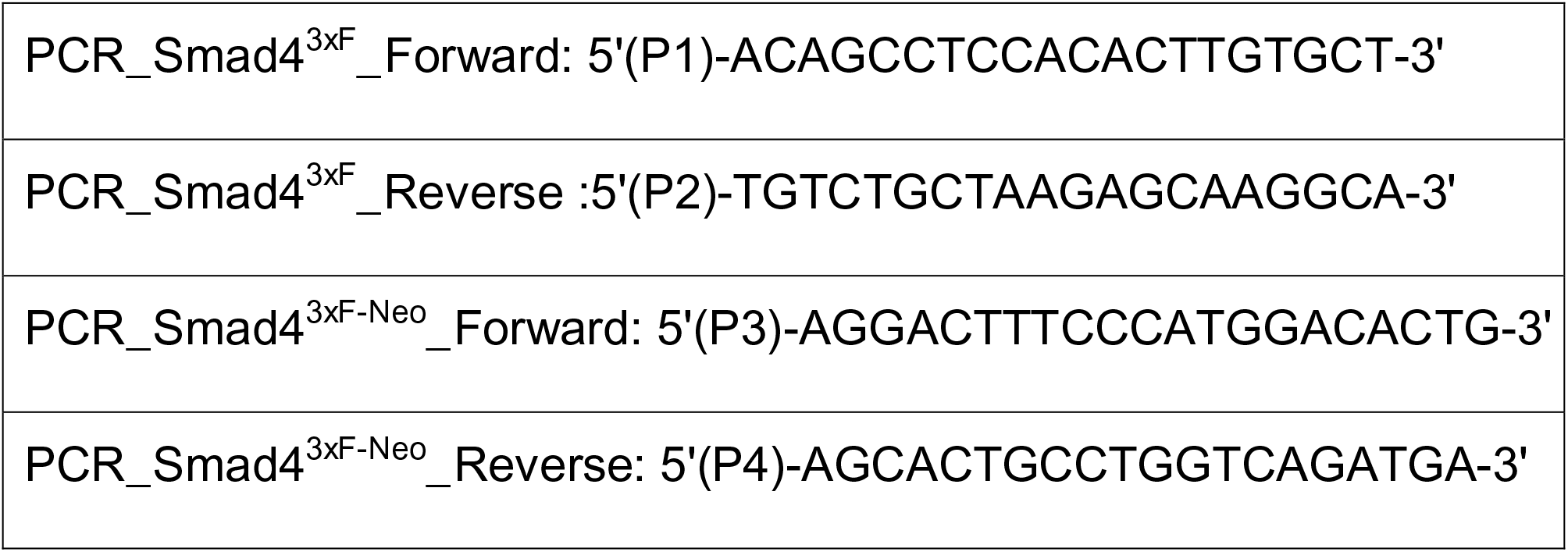

### SMAD4-3xFlag ChIP-seq

About 80 *Smad4^3xF/3xF^* embryos at E9.5-E10.0 (25 to 30 somites; forelimbs with proximal trunk tissues) or 100 *Smad4^3xF/3xF^* embryos E10.5 (forelimbs/hindlimbs) were dissected per replicate. The ChIP protocol was performed as previously described (Osterwalder et al., 2014) with one modification: SMAD4-chromatin complexes were immuno-precipitated only for 6 hours instead of overnight to reduce non-specific background. Libraries for sequencing were constructed using the KAPA Hyper Prep Kit (ref KK8502) and sequenced using the Illumina NextSeq 500 system.

### ChIP-seq analysis and annotation

Short reads obtained from Illumina NextSeq were aligned to the mm9 genome using Bowtie v1.1.0 (Langmead et al., 2009). Only those reads with a unique match to the genome with two or fewer mismatches (*-m 1 -v 2*) were retained. In order to make different runs comparable, the 3’ of reads were trimmed to 63 bp before alignment. This step was performed using fastx_trimmer (*-l 63*), a tool part of the FASTX-Toolkit (http://hannonlab.cshl.edu/fastx_toolkit/) (v0.0.13). Peak calling was performed using MACS v1.4 (Zhang et al., 2008) with the following parameters: *--gsize=mm --bw=300 --nomodel --shiftsize=100 --pvalue=1e-2*. Input DNA from the same sample was used as a control. Wiggle tracks were also generated with MACS, these were then re-scaled linearly according to sequencing depth (RPM, Reads Per Million sequenced reads). MACS was run with a permissive threshold (*p*-value 0.01) in order to identify a larger list of sub-significant regions across biological replicates. Evidences from these replicates were combined using MSPC (Jalili et al., 2015), with the following parameters *-r biological -s 1E-5 -W 1E-2*. The confirmed peaks were assigned the best *p*-value (as defined by MACS) among the overlapping peaks across replicates. Only replicated peaks were retained for further analysis (termed as *golden* for convenience; one golden set per developmental stage). These lists of peaks were annotated to the TSS of the nearest RefSeq genes using the script *annotatePeaks.pl* available in HOMER (Heinz et al., 2010). A region was considered as proximal to a promoter if located within 2.5 kb from a RefSeq promoter. The remaining regions were divided into intragenic and intergenic, whether the region overlapped the body of an annotated gene or not.

### Motif enrichment and *de novo* motif discovery analyses

The script *findMotifsGenome.pl* available in HOMER (Heinz et al., 2010) was used to perform enrichment analysis for known transcription-factor binding sites and motif discovered *de novo*. The script was run with the following arguments: *-size -150,150 - len 6,7,8,9,10,12,14*, using the peak summits of the *golden* set as reference. The top ten most significant, over-represented known matrices along with the top ten *de novo* discovered motifs were then used to scan every single region for high-affinity sites using FIMO (v4.10.0; Grant et al., 2011). The following parameters were used: *-- thresh 1e-4 --no-qvalue*. The resulting list of sites was transformed into a matrix in which each region was represented as a vector of *p*-values, one for each different motif, corresponding to the *p*-value of the highest-scoring site identified (*p*-value = 1 if no significant match was found). *P*-values were then log10-transformed and their sign inverted, then hierarchically clustered (*hclust* function of R; Euclidean distance; complete linkage).

### Evolutionary conservation analysis of the genomic regions enriched in SMAD4-chromatin complexes

The genome-wide track of base-pair *Phastcons* (Siepel et al., 2005) conservation scores in placental mammals was downloaded from the UCSC genome browser (Tyner et al., 2017) (track name: *mm10.60way.phastCons60wayPlacental.bw*). The coordinates of the peaks in the golden sets were converted from mm9 to mm10 using *liftOver* (Tyner et al., 2017) (*-minMatch=0.95*). The base-pair scores for the 300 bp centered on the summit of the peaks were then extracted using *bwtool* (Pohl and Beato, 2014).

### ChIP-qPCR analysis

Two BMP responsive elements (BRE of Msx2 and BRE of Id1; Brugger et al., 2004; Korchynskyi and ten Dijke, 2002) identified in the ChIP-seq dataset were validated by ChIP-qPCR. Each duplicate contains 45 pairs of forelimbs with proximal trunk tissues from *Smad4^3xF/3xF^* embryos at E9.75 or fore- and hindlimbs buds at E10.5. An unlinked amplicon within the β*-actin* locus was used as normalizing control and to calculate the fold-enrichment. A qPCR cycle threshold of 32 was defined as background enrichment. For each experiment, two genomic regions not enriched in the SMAD4^3xF^ ChIP-seq dataset were used as negative controls. These are oligos used for qPCR amplification:

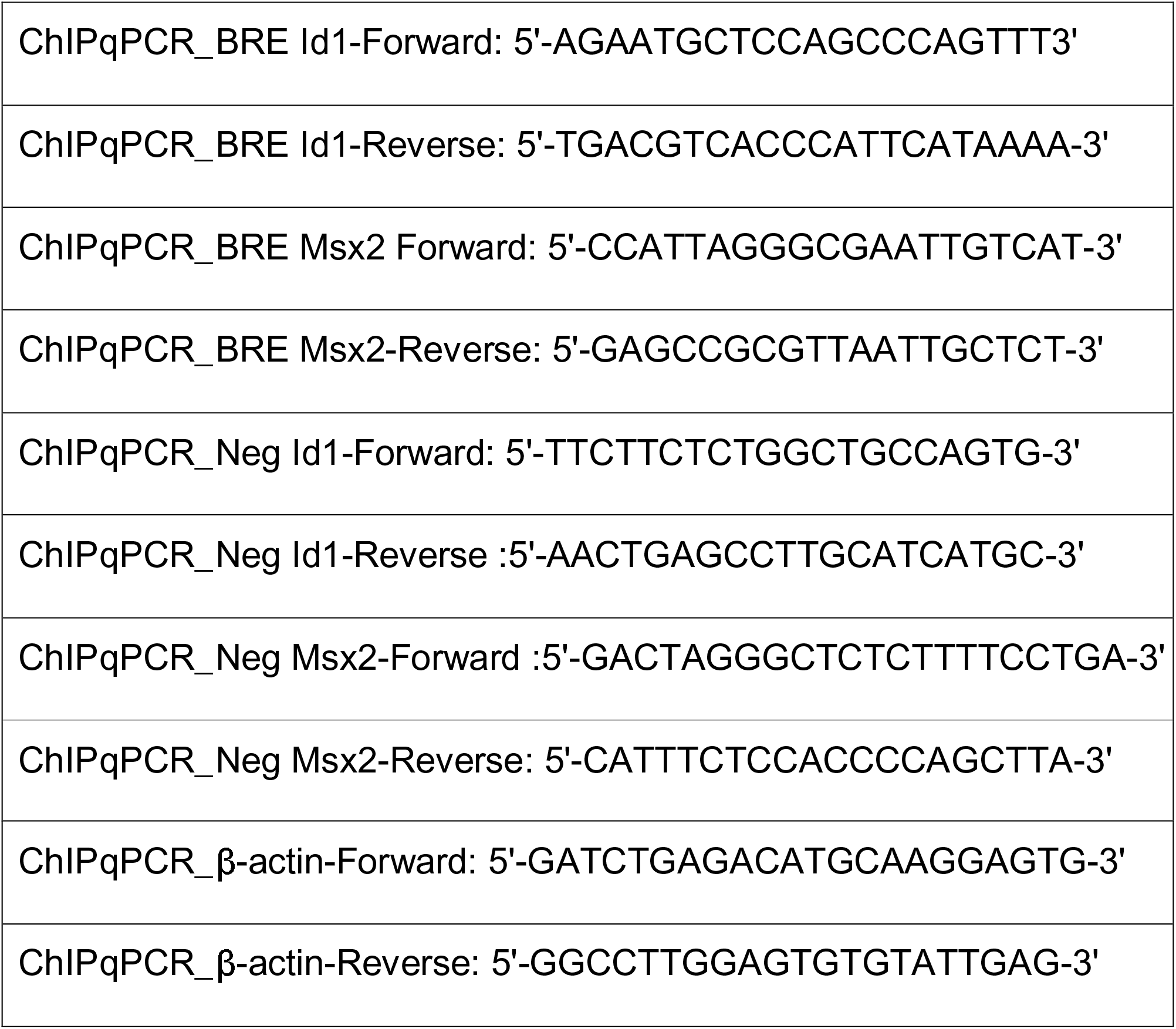

### ATAC-seq analysis and annotation

For the early stage, each replicate contains a pair of forelimbs with proximal trunk tissues isolated from wild-type embryos at E9.75. For the later stage, each replicate consists of a pair of forelimbs isolated from wild-type embryo at E10.5. For both stages, two biological replicates were processed independently as described (Buenrostro et al., 2013). The ATAC libraries were prepared by amplifying the transposed DNA fragments with the KAPA HiFI HotStart ReadyMix kit followed by sequencing on an Illumina NextSeq 500. The short reads were aligned to the mm9 genome using Bowtie v1.1.0 (Langmead et al., 2009; -m 1 -v 2, see “Chip-seq raw data analyses and annotation”). Accessible regions were identified using MACS v1.4 (Zhang et al., 2008) with the following parameters: *--gsize=mm --bw=150 --nomodel - -nolambda --shiftsize=75*. Genome-wide profiles were generated using MACS and re-scaled linearly according to sequencing depth (RPM). Gene annotation was performed using HOMER (Heinz et al., 2010), as described in *“Chip-seq raw data analyses and annotation”*. Evidence from biological replicates was combined using MSPC (Jalili et al., 2015), using the following parameters *-r biological -s 1E-10 -W 1E-6*. The confirmed regions were assigned the best *p*-value (as defined by MACS) among the overlapping regions across replicates.

### RNA-seq analysis

Dissected wild-type and *Smad4*^Δ/Δ^ forelimb buds from E10.0 embryos (30 somites) and 10.5 embryos (35 somites) were collected in RNAlater® (Sigma R0901), incubated overnight at 4°C and then stored at -80°C. Both forelimb buds of one biological replicate were pooled. After genotyping, 4 age-matched and gender-matched *Smad4*^Δ/Δ^ forelimb bud pairs and 3 wild-type replicates per stage were sequenced. RNA was extracted using the Qiagen RNeasy micro kit. For each replicate, the quality of total RNA was analysed using the RNA 6000 Pico kit (Agilent 2100 bioanalyzer), which was followed by polyA-mediated RNA library preparation. Sequencing was done one a HiSeq 2500 machine using the single-read 50 cycles protocol.

Single-end reads obtained from Illumina HiSeq were aligned to the mm9 reference genome and to the *Mus Musculus* transcriptome (iGenome refGene GTF) using TopHat v2.0.13 (Kim et al., 2013). The option *--no-coverage-search* was specified, while all the other parameters were left to default. Only uniquely mapped reads were considered for the analysis. Tracks for the UCSC genome browser (Tyner et al., 2017) were produced using *genomeCoverageBed* from BedTools v2.17.0 (Quinlan and Hall, 2010); these were linearly re-scaled according to sequencing depth (RPM). Gene-wise counts were computed using *htseq-count* from the HTSeq package (Anders et al., 2015) with *-s* set to *no*. Genes on chromosomes X, Y and M were excluded from further analysis. edgeR (Robinson et al., 2010) was used to identify differentially expressed genes (DEGs). Only genes showing expression (in terms of Fragments Per Million sequenced reads equal or higher than 1) in at least three samples were considered for further analyses. Libraries were normalized according to TMM normalization. Tag-wise estimation of dispersion was evaluated using *prior.df = 10*. Differential expression between pairs of conditions was evaluated using the *exactTest* R function. False discovery rates were estimated using Benjamini-Hochberg correction (Benjamini and Hochberg, 1995). DEGs were defined as those genes showing a *q*-value <= 0.1 and a linear fold-change equal or higher than 1.2. Functional enrichment analyses were conducted using DAVID (Huang da et al., 2009).

The SMAD4-bound regions were associated with their target genes using the topologically associating domains (TADs) defined in mouse ES cells (Dixon et al., 2012). At a particular developmental stage, the expressed genes were classified as either unchanged, up- or down-regulated. Each gene was assigned to the corresponding TAD, and the number of SMAD4-binding peaks per TAD was calculated and normalized to the total number of genes within in the domain. Using this strategy, it was possible to assign the SMAD4-interacting genomic regions to particular genes.

### Hierarchical clustering, plots and statistical testing

Clustering, plots, heat maps and statistics were handled in the statistical computing environment R v3.

### Immunofluorescence analysis

Embryos were collected in ice-cold PBS and fixed for 2 hrs at 4°C in 4% PFA/PBS. Samples were then cryoprotected using a gradient of sucrose: 10% sucrose/PBS (w/v), 20% sucrose/PBS, 30% sucrose/PBS (1hr each) at 4°C. Embryos were then embedded 50:50 (v/v) OCT/30% sucrose. For immunofluorescent staining, 10µm sections were prepared. *Smad4^3xF/3xF^* or wild-type sections were washed 3×5min in PBS, once 30min in PBT and again 5min in PBS. They were blocked in 1% BSA in PBT for 1hr at RT and incubated overnight at 4°C with the monoclonal mouse anti-FLAG M2 antibody (Sigma, F1804) diluted 1:500 in 1% BSA/PBS. Sections were washed 3×5min in PBS, once in PBT and were incubated in the dark for 1hr at RT with the goat anti-mouse Alexa 488 secondary antibody diluted 1:500 in 1% BSA/PBS. Sections were finally washed 3×10min PBS, once in PBT (5min), nuclei counterstained in 1µg/mL Hoechst-33258/PBS (5min) and rinsed again 3×5min in PBS. Then they were mounted in Mowiol 4-88 and dried overnight at RT in the dark.

### Whole Mount *in situ* hybridization (WISH)

Embryos were age-matched by counting somites. Whole-mount *in situ* hybridization was performed using standard protocols. *Smad4^+/+^ Prx1-Cre^Tg/+^* embryos were always used as wild-type controls (WT).

### Generation and analysis of transgenic *LacZ* founder embryos

Candidate CRM regions were amplified by PCR from mouse genomic DNA and were then cloned into a Hsp68-*LacZ* reporter vector (Osterwalder et al., 2014) using the Gibson Assembly® Method. Transgenic embryos were generated by pronuclear injection.

### Culture of wild-type and mutant LMPs in cholesterol-free medium and quantitation of intracellular cholesterol

Forelimb buds (E10.0, 28-29 somites) were collected into ice-cold PBS and incubated in cold 2% Trypsin (Gibco 15090-046)/PBS at 4°C for 30min. The reaction was stopped by adding an excess of DMEM medium containing 10% fetal bovine serum (FBS). The limb bud ectoderm was then removed. By gently pipetting, LMPs were dissociated and seeded in 2-3 wells of a 96 well plate with DMEM medium containing 10% lipid-depleted FBS (PanBiotech P30.3302), 4.5g/L Glucose (Gibco 41966-029), 100U Penicillin, 0.1mg/mL Streptomycin (Sigma P-0781) and 200mM L-Glutamine (Sigma G-7513). LMPs were cultured in this cholesterol-free medium for 20 t0 24 hrs and then treated with 10mM methyl-β-cyclodextrin (MβCD, Sigma C4555) for 15min at 37°C to remove cholesterol from plasma membranes. Then, LMPs were trypsinized gently for 2min in 2% trypsin and LMPs from 2 pairs of forelimb buds of the same genotype pooled for one biological replicate. After determining cell numbers, LMPs were mixed with 2mL of ethanol/chloroform solution (2:1) in a glass tube. Following 5 min centrifugation at 1400g, the supernatant was transferred into a new glass tube and mixed with 250µL 50mM citric acid, 500µL water and 250µL chloroform. After 30min centrifugation at 1400g, the lowest phase-containing lipids was transferred in a 1.5mL Eppendorf tube and dried using a Speedvac centrifuge. The dried lipids were solubilized in 95% ethanol and the cholesterol was quantitated using the Amplex Red Cholesterol Assay kit (Invitrogen A12216). All values for the wild-type and mutant samples are shown in Table S9. Please note that all measured values for the wild-type samples were clearly above the detection limit. The concentrations of intracellular cholesterol were calculated by dividing the measured cholesterol levels by the cell numbers determined.

Preparation of lipid/cholesterol depleted fetal bovine serum (FBS): 500mL FBS were stirred overnight at 4°C with 10g Cab-osil M-5 (ACROS Organics 7631-86-9). The mix was then centrifuged for 10min at 3000rpm and the supernatant filtrated under sterile conditions. The Insulin-Transferrin-Sodium-Selenite media supplement (Sigma I-1884-1) was dissolved in 50ml of H_2_O, acidified by adding 250µL HCl and filtrated under sterile conditions. 25mL of the Insulin-Transferrin-Sodium-Selenite solution was added to 500mL of lipid-depleted FCS. Aliquots of 30mL were frozen at -20°C and used for preparing 500 mL of lipid-depleted EMFI medium.

### Imaging

Images were taken using a Leica MZ FLII stereomicroscope and the Leica Application Suite V3 software. Contrast and image sized were adjusted with Adobe Photoshop CS5.1. All bar-plots were generated using GraphPad Prism 7. All figures were generated with Adobe Photoshop or Adobe Illustrator.

### Statistical Analysis

#### ChIP-seq

After sequence alignment, peak calling was performed using MACS v1.4 (Zhang et al., 2008) with a *p-value threshold of 1e-2*.

#### RNA-seq

Following the alignment of sequences, edgeR (Robinson et al., 2010) was used to identify differentially expressed genes (DEGs). DEGs were defined as those genes showing an FDR ≤ 0.1 and an absolute linear fold-change equal or higher than 1.2.

### Data availability

All ChIP-seq, ATAC-seq and RNA-seq datasets have been deposited in the Gene Expression Omnibus (GEO) database under the accession number GSE114257 [https://www.ncbi.nlm.nih.gov/geo/query/acc.cgi?acc=GSE114257].

## Acknowledgements

We thank J. Stolte for technical assistance, F. Gullotta for help with LMP culture and J. Lopez-Rios for sharing ATAC-seq datasets. J. Malkmus is thanked for graciously providing the *Shh^−/-^* panels in Fig. S6. We are grateful to A. Offinger’s team for excellent animal care, P. Pelczar and his team (Centre for Transgenic Models at the University of Basel) for generating the *LacZ* founder embryos and P. Lorentz (DBM Bio-Optics Core Facility) for imaging support. This research was initially supported by SNF grant 310030B_166685 and then the ERC-2015-AdG Project INTEGRAL (ID 695032). IB and AV were supported by NIH grants R01HG003988, U54HG006997 and UM1HG009421. Research at the E.O. Lawrence Berkeley National Laboratory was performed under Dept. of Energy Contract DE-AC02-05CH11231 to University of California.

## Author contributions

JG performed most of the experiments in this study. IB performed all the bioinformatics analysis as a postdoctoral fellow in the research group of AV and more recently as an independent investigator. FL generated the *Smad4*^3xF^ allele and performed the initial analysis of the *Smad4*^3xF^ mice. FL and JG established the methods for sensitive and specific detection of the endogenous SMAD4^3xF^ protein and its suitability for ChIP analysis. RR, LRM and TO contributed to the whole mount RNA *in situ* analysis of forelimb buds. AZ performed the GRN analysis shown in Fig. 7 with input from IB. RZ and AZ conceived and supervised the study, acquired the necessary funding and wrote the manuscript with input from all authors.

## Supplementary information

Supplemental Information includes Figures S1-S6 and Tables S1-S15.

## References

Ahn, K., Mishina, Y., Hanks, M. C., Behringer, R. R. and Crenshaw, E. B., 3rd (2001). BMPR-IA signaling is required for the formation of the apical ectodermal ridge and dorsal-ventral patterning of the limb. Development 128, 4449–4461.

Anders, S., Pyl, P. T. and Huber, W. (2015). HTSeq--a Python framework to work with high-throughput sequencing data. Bioinformatics 31, 166–169.

Bailey, J. S., Rave-Harel, N., McGillivray, S. M., Coss, D. and Mellon, P. L. (2004). Activin regulation of the follicle-stimulating hormone beta-subunit gene involves Smads and the TALE homeodomain proteins Pbx1 and Prep1. Molecular endocrinology 18, 1158–1170.

Bandyopadhyay, A., Tsuji, K., Cox, K., Harfe, B. D., Rosen, V. and Tabin, C. J. (2006). Genetic Analysis of the Roles of BMP2, BMP4, and BMP7 in Limb Patterning and Skeletogenesis. Plos Genet 2, e216.

Bastida, M. F. l., Sheth, R. and Ros, M. A. (2009). A BMP-Shh negative-feedback loop restricts Shh expression during limb development. Development 136, 3779–3789.

Bénazet, J.-D., Bischofberger, M., Tiecke, E., Gonçalves, A., Martin, J. F., Zuniga, A., Naef, F. and Zeller, R. (2009). A Self-Regulatory System of Interlinked Signaling Feedback Loops Controls Mouse Limb Patterning. Science 323, 1050–1053.

Bénazet, J.-D., Pignatti, E., Nugent, A., Unal, E., Laurent, F. and Zeller, R. (2012). Smad4 is required to induce digit ray primordia and to initiate the aggregation and differentiation of chondrogenic progenitors in mouse limb buds. Development 139, 4250–4260.

Bénazet, J. D. and Zeller, R. (2013). Dual requirement of ectodermal Smad4 during AER formation and termination of feedback signaling in mouse limb buds. Genesis 51, 660–666.

Benjamini, Y. and Hochberg, Y. (1995). Controlling the False Discovery Rate - a Practical and Powerful Approach to Multiple Testing. J Roy Stat Soc B Met 57, 289–300.

Berge, D. t., Brouwer, A., Korving, J., Martin, J. F. and Meijlink, F. (1998). Prx1 and Prx2 in skeletogenesis: roles in the craniofacial region, inner ear and limbs. Development (Cambridge, England) 125, 3831–3842.

Bitgood, M. J. and McMahon, A. P. (1995). Hedgehog and Bmp genes are coexpressed at many diverse sites of cell-cell interaction in the mouse embryo. Dev Biol 172, 126–138.

Briscoe, J. and Therond, P. P. (2013). The mechanisms of Hedgehog signalling and its roles in development and disease. Nature reviews. Molecular cell biology 14, 416–429.

Brugger, S. M., Merrill, A. E., Torres-Vazquez, J., Wu, N., Ting, M. C., Cho, J. Y., Dobias, S. L., Yi, S. E., Lyons, K., Bell, J. R., et al. (2004). A phylogenetically conserved cis-regulatory module in the Msx2 promoter is sufficient for BMP-dependent transcription in murine and Drosophila embryos. Development 131, 5153–5165.

Buenrostro, J. D., Giresi, P. G., Zaba, L. C., Chang, H. Y. and Greenleaf, W. J. (2013). Transposition of native chromatin for fast and sensitive epigenomic profiling of open chromatin, DNA-binding proteins and nucleosome position. Nature methods 10, 1213–1218.

Chen, Y. and Gridley, T. (2013). Compensatory regulation of the Snai1 and Snai2 genes during chondrogenesis. Journal of bone and mineral research : the official journal of the American Society for Bone and Mineral Research 28, 1412–1421.

Chu, G. C., Dunn, N. R., Anderson, D. C., Oxburgh, L. and Robertson, E. J. (2004). Differential requirements for Smad4 in TGFbeta-dependent patterning of the early mouse embryo. Development 131, 3501–3512.

Chuang, P. T., Kawcak, T. and McMahon, A. P. (2003). Feedback control of mammalian Hedgehog signaling by the Hedgehog-binding protein, Hip1, modulates Fgf signaling during branching morphogenesis of the lung. Genes Dev 17, 342–347.

Chuang, P. T. and McMahon, A. P. (1999). Vertebrate Hedgehog signalling modulated by induction of a Hedgehog-binding protein. Nature 397, 617–621.

Cooper, M. K., Wassif, C. A., Krakowiak, P. A., Taipale, J., Gong, R., Kelley, R. I., Porter, F. D. and Beachy, P. A. (2003). A defective response to Hedgehog signaling in disorders of cholesterol biosynthesis. Nat Genet 33, 508–513.

Dixon, J. R., Selvaraj, S., Yue, F., Kim, A., Li, Y., Shen, Y., Hu, M., Liu, J. S. and Ren, B. (2012). Topological domains in mammalian genomes identified by analysis of chromatin interactions. Nature 485, 376–380.

Dunker, N., Schmitt, K. and Krieglstein, K. (2002). TGF-beta is required for programmed cell death in interdigital webs of the developing mouse limb. Mech Dev 113, 111–120.

Farin, H. F., Ludtke, T. H., Schmidt, M. K., Placzko, S., Schuster-Gossler, K., Petry, M., Christoffels, V. M. and Kispert, A. (2013). Tbx2 terminates shh/fgf signaling in the developing mouse limb bud by direct repression of gremlin1. Plos Genet 9, e1003467.

Geetha-Loganathan, P., Nimmagadda, S. and Scaal, M. (2008). Wnt signaling in limb organogenesis. Organogenesis 4, 109–115.

Grant, C. E., Bailey, T. L. and Noble, W. S. (2011). FIMO: scanning for occurrences of a given motif. Bioinformatics 27, 1017–1018.

Harfe, B. D., Scherz, P. J., Nissim, S., Tian, H., McMahon, A. P. and Tabin, C. J. (2004). Evidence for an expansion-based temporal Shh gradient in specifying vertebrate digit identities. Cell 118, 517–528.

Harris, M. P., Fallon, J. F. and Prum, R. O. (2002). Shh-Bmp2 signaling module and the evolutionary origin and diversification of feathers. J Exp Zool 294, 160–176.

Heinz, S., Benner, C., Spann, N., Bertolino, E., Lin, Y. C., Laslo, P., Cheng, J. X., Murre, C., Singh, H. and Glass, C. K. (2010). Simple combinations of lineage-determining transcription factors prime cis-regulatory elements required for macrophage and B cell identities. Mol Cell 38, 576–589.

Horvat, S., McWhir, J. and Rozman, D. (2011). Defects in cholesterol synthesis genes in mouse and in humans: lessons for drug development and safer treatments. Drug Metab Rev 43, 69–90.

Hsu, D. R., Economides, A. N., Wang, X., Eimon, P. M. and Harland, R. M. (1998). The Xenopus dorsalizing factor Gremlin identifies a novel family of secreted proteins that antagonize BMP activities. Mol Cell 1, 673–683.

Huang da, W., Sherman, B. T. and Lempicki, R. A. (2009). Systematic and integrative analysis of large gene lists using DAVID bioinformatics resources. Nat Protoc 4, 44–57.

Imuta, Y., Nishioka, N., Kiyonari, H. and Sasaki, H. (2009). Short limbs, cleft palate, and delayed formation of flat proliferative chondrocytes in mice with targeted disruption of a putative protein kinase gene, Pkdcc (AW548124). Dev Dyn 238, 210–222.

Jalili, V., Matteucci, M., Masseroli, M. and Morelli, M. J. (2015). Using combined evidence from replicates to evaluate ChIP-seq peaks. Bioinformatics 31, 2761–2769.

Karamboulas, K., Dranse, H. J. and Underhill, T. M. (2010). Regulation of BMP-dependent chondrogenesis in early limb mesenchyme by TGFbeta signals. Journal of cell science 123, 2068–2076.

Karaplis, A. C., Luz, A., Glowacki, J., Bronson, R. T., Tybulewicz, V. L., Kronenberg, H. M. and Mulligan, R. C. (1994). Lethal skeletal dysplasia from targeted disruption of the parathyroid hormone-related peptide gene. Genes Dev 8, 277–289.

Katagiri, T., Boorla, S., Frendo, J. L., Hogan, B. L. M. and Karsenty, G. (1998). Skeletal abnormalities in doubly heterozygous Bmp4 and Bmp7 mice. Dev Genet 22, 340–348.

Kim, D., Pertea, G., Trapnell, C., Pimentel, H., Kelley, R. and Salzberg, S. L. (2013). TopHat2: accurate alignment of transcriptomes in the presence of insertions, deletions and gene fusions. Genome Biol 14, R36.

Korchynskyi, O. and ten Dijke, P. (2002). Identification and functional characterization of distinct critically important bone morphogenetic protein-specific response elements in the Id1 promoter. The Journal of biological chemistry 277, 4883–4891.

Krakowiak, P. A., Wassif, C. A., Kratz, L., Cozma, D., Kovarova, M., Harris, G., Grinberg, A., Yang, Y., Hunter, A. G., Tsokos, M., et al. (2003).

Lathosterolosis: an inborn error of human and murine cholesterol synthesis due to lathosterol 5-desaturase deficiency. Hum Mol Genet 12, 1631–1641.

Kuijper, S., Feitsma, H., Sheth, R., Korving, J., Reijnen, M. and Meijlink, F. (2005). Function and regulation of Alx4 in limb development: complex genetic interactions with Gli3 and Shh. Dev Biol 285, 533–544.

Lallemand, Y., Nicola, M.-A., Ramos, C., Bach, A., Cloment, C. c. S. and Robert, B. t. (2005). Analysis of Msx1; Msx2 double mutants reveals multiple roles for Msx genes in limb development. Development 132, 3003–3014.

Langmead, B., Trapnell, C., Pop, M. and Salzberg, S. L. (2009). Ultrafast and memory-efficient alignment of short DNA sequences to the human genome. Genome Biol 10, R25.

Li, J., Feng, J., Liu, Y., Ho, T. V., Grimes, W., Ho, H. A., Park, S., Wang, S. and Chai, Y. (2015). BMP-SHH signaling network controls epithelial stem cell fate via regulation of its niche in the developing tooth. Dev Cell 33, 125–135.

Li, Y., Zhang, H., Litingtung, Y. and Chiang, C. (2006). Cholesterol modification restricts the spread of Shh gradient in the limb bud. Proc Natl Acad Sci U S A 103, 6548–6553.

Lim, J., Tu, X., Choi, K., Akiyama, H., Mishina, Y. and Long, F. (2015). BMP– Smad4 signaling is required for precartilaginous mesenchymal condensation independent of Sox9 in the mouse. Dev Biol 400, 132–138.

Logan, M., Martin, J. F., Nagy, A., Lobe, C., Olson, E. N. and Tabin, C. J. (2002). Expression of Cre recombinase in the developing mouse limb bud driven by a Prxl enhancer. Genesis 33, 77–80.

Lopez-Rios, J., Speziale, D., Robay, D., Scotti, M., Osterwalder, M., Nusspaumer, G., Galli, A., Holländer, Georg A., Kmita, M. and Zeller, R. (2012). GLI3 Constrains Digit Number by Controlling Both Progenitor Proliferation and BMP-Dependent Exit to Chondrogenesis. Dev Cell 22, 837–848.

Lopez-Rovira, T., Chalaux, E., Massague, J., Rosa, J. L. and Ventura, F. (2002). Direct binding of Smad1 and Smad4 to two distinct motifs mediates bone morphogenetic protein-specific transcriptional activation of Id1 gene. The Journal of biological chemistry 277, 3176–3185.

Lorda-Diez, C. I., Duarte-Olivenza, C., Hurle, J. M. and Montero, J. A. (2021). Transforming growth factor beta signaling: The master sculptor of fingers. Dev Dyn.

Luo, G., Hofmann, C., Bronckers, A. L., Sohocki, M., Bradley, A. and Karsenty, G. (1995). BMP-7 is an inducer of nephrogenesis, and is also required for eye development and skeletal patterning. Genes Dev 9, 2808–2820.

Luo, J., Yang, H. and Song, B. L. (2020). Mechanisms and regulation of cholesterol homeostasis. Nature reviews. Molecular cell biology 21, 225–245.

Madison, B. B., Braunstein, K., Kuizon, E., Portman, K., Qiao, X. T. and Gumucio, D. L. (2005). Epithelial hedgehog signals pattern the intestinal crypt-villus axis. Development 132, 279–289.

Malkmus, J., Ramos Martins, L., Jhanwar, S., Kircher, B., Palacio, V., Sheth, R., Leal, F., Duchesne, A., Lopez-Rios, J., Peterson, K. A., et al. (in press). Spatial cooperativity among multiple *Gremlin1* enhancers provides digit development with cis-regulatory robustness and evolutionary plasticity. Nat Commun.

Mateus, R., Holtzer, L., Seum, C., Hadjivasiliou, Z., Dubois, M., Julicher, F. and Gonzalez-Gaitan, M. (2020). BMP Signaling Gradient Scaling in the Zebrafish Pectoral Fin. Cell Rep 30, 4292–4302 e4297.

Mukhopadhyay, M., Shtrom, S., Rodriguez-Esteban, C., Chen, L., Tsukui, T., Gomer, L., Dorward, D. W., Glinka, A., Grinberg, A., Huang, S.-P., et al. (2001). Dickkopf1 Is Required for Embryonic Head Induction and Limb Morphogenesis in the Mouse. Dev Cell 1, 423–434.

Norrie, J. L., Lewandowski, J. P., Bouldin, C. M., Amarnath, S., Li, Q., Vokes, M. S., Ehrlich, L. I. R., Harfe, B. D. and Vokes, S. A. (2014). Dynamics of BMP signaling in limb bud mesenchyme and polydactyly. Dev Biol 393, 270–281.

Okada, I., Hamanoue, H., Terada, K., Tohma, T., Megarbane, A., Chouery, E., Abou-Ghoch, J., Jalkh, N., Cogulu, O., Ozkinay, F., et al. (2011). SMOC1 Is Essential for Ocular and Limb Development in Humans and Mice. Am J Hum Genetics 88, 30–41.

Osterwalder, M., Speziale, D., Shoukry, M., Mohan, R., Ivanek, R., Kohler, M., Beisel, C., Wen, X., Scales, S. J., Christoffels, V. M., et al. (2014). HAND2 targets define a network of transcriptional regulators that compartmentalize the early limb bud mesenchyme. Dev Cell 31, 345–357.

Ovchinnikov, D. A., Selever, J., Wang, Y., Chen, Y.-T., Mishina, Y., Martin, J. F. and Behringer, R. R. (2006). BMP receptor type IA in limb bud mesenchyme regulates distal outgrowth and patterning. Dev Biol 295, 103–115.

Pajni-Underwood, S., Wilson, C. P., Elder, C., Mishina, Y. and Lewandoski, M. (2007). BMP signals control limb bud interdigital programmed cell death by regulating FGF signaling. Development 134, 2359–2368.

Panman, L., Galli, A., Lagarde, N., Michos, O., Soete, G., Zuniga, A. and Zeller, R. (2006). Differential regulation of gene expression in the digit forming area of the mouse limb bud by SHH and gremlin 1/FGF-mediated epithelial-mesenchymal signalling. Development 133, 3419–3428.

Patten, I. and Placzek, M. (2002). Opponent activities of Shh and BMP signaling during floor plate induction in vivo. Curr Biol 12, 47–52.

Pelikan, R. C., Iwata, J., Suzuki, A., Chai, Y. and Hacia, J. G. (2013). Identification of candidate downstream targets of TGFbeta signaling during palate development by genome-wide transcript profiling. J Cell Biochem 114, 796–807.

Pignatti, E., Zeller, R. and Zuniga, A. (2014). To BMP or not to BMP during vertebrate limb bud development. Semin Cell Dev Biol 32, 119–127.

Pizette, S. and Niswander, L. (2000). BMPs are required at two steps of limb chondrogenesis: formation of prechondrogenic condensations and their differentiation into chondrocytes. Dev Biol 219, 237–249.

Pohl, A. and Beato, M. (2014). bwtool: a tool for bigWig files. Bioinformatics 30, 1618–1619.

Porter, F. D. and Herman, G. E. (2011). Malformation syndromes caused by disorders of cholesterol synthesis. Journal of lipid research 52, 6–34.

Probst, S., Kraemer, C., Demougin, P., Sheth, R., Martin, G. R., Shiratori, H., Hamada, H., Iber, D., Zeller, R. and Zuniga, A. (2011). SHH propagates distal limb bud development by enhancing CYP26B1-mediated retinoic acid clearance via AER-FGF signalling. Development 138, 1913–1923.

Probst, S., Zeller, R. and Zuniga, A. (2013). The hedgehog target Vlk genetically interacts with Gli3 to regulate chondrocyte differentiation during mouse long bone development. Differentiation 85, 121–130.

Quinlan, A. R. and Hall, I. M. (2010). BEDTools: a flexible suite of utilities for comparing genomic features. Bioinformatics 26, 841–842.

Radhakrishnan, A., Rohatgi, R. and Siebold, C. (2020). Cholesterol access in cellular membranes controls Hedgehog signaling. Nat Chem Biol 16, 1303–1313.

Reinhardt, R., Gullotta, F., Nusspaumer, G., Ünal, E., Ivanek, R., Zuniga, A. and Zeller, R. (2019). Molecular signatures identify immature mesenchymal progenitors in early mouse limb buds that respond differentially to morphogen signaling. Development 146, dev.173328.

Robinson, M. D., McCarthy, D. J. and Smyth, G. K. (2010). edgeR: a Bioconductor package for differential expression analysis of digital gene expression data. Bioinformatics 26, 139–140.

Scherz, P. J., Harfe, B. D., McMahon, A. P. and Tabin, C. J. (2004). The limb bud Shh-Fgf feedback loop is terminated by expansion of former ZPA cells. Science 305, 396–399.

Selever, J., Liu, W., Lu, M.-F., Behringer, R. R. and Martin, J. F. (2004). Bmp4 in limb bud mesoderm regulates digit pattern by controlling AER development. Dev Biol 276, 268–279.

Siepel, A., Bejerano, G., Pedersen, J. S., Hinrichs, A. S., Hou, M., Rosenbloom, K., Clawson, H., Spieth, J., Hillier, L. W., Richards, S., et al. (2005). Evolutionarily conserved elements in vertebrate, insect, worm, and yeast genomes. Genome Res 15, 1034–1050.

Taher, L., Collette, N. M., Murugesh, D., Maxwell, E., Ovcharenko, I. and Loots, G. G. (2011). Global gene expression analysis of murine limb development. Plos One 6, e28358.

Thomas, J. T., Eric Dollins, D., Andrykovich, K. R., Chu, T., Stultz, B. G., Hursh, D. A. and Moos, M. (2017). SMOC can act as both an antagonist and an expander of BMP signaling. Elife 6.

Tint, G. S., Yu, H., Shang, Q., Xu, G. and Patel, S. B. (2006). The use of the Dhcr7 knockout mouse to accurately determine the origin of fetal sterols. Journal of lipid research 47, 1535–1541.

Tissieres, V., Geier, F., Kessler, B., Wolf, E., Zeller, R. and Lopez-Rios, J. (2020). Gene Regulatory and Expression Differences between Mouse and Pig Limb Buds Provide Insights into the Evolutionary Emergence of Artiodactyl Traits. Cell Rep 31, 107490.

Tyner, C., Barber, G. P., Casper, J., Clawson, H., Diekhans, M., Eisenhart, C., Fischer, C. M., Gibson, D., Gonzalez, J. N., Guruvadoo, L., et al. (2017). The UCSC Genome Browser database: 2017 update. Nucleic Acids Res 45, D626–D634.

Tzchori, I., Day, T. F., Carolan, P. J., Zhao, Y., Wassif, C. A., Li, L., Lewandoski, M., Gorivodsky, M., Love, P. E., Porter, F. D., et al. (2009). LIM homeobox transcription factors integrate signaling events that control three-dimensional limb patterning and growth. Development 136, 1375–1385.

Verheyden, J. M. and Sun, X. (2008). An Fgf/Gremlin inhibitory feedback loop triggers termination of limb bud outgrowth. Nature 454, 638–641.

Vienken, H., Mabrouki, N., Grabau, K., Claas, R. F., Rudowski, A., Schomel, N., Pfeilschifter, J., Lutjohann, D., van Echten-Deckert, G. and Meyer Zu Heringdorf, D. (2017). Characterization of cholesterol homeostasis in sphingosine-1-phosphate lyase-deficient fibroblasts reveals a Niemann-Pick disease type C-like phenotype with enhanced lysosomal Ca(2+) storage. Sci Rep 7, 43575.

Weiss, A. and Attisano, L. (2013). The TGFbeta superfamily signaling pathway. Wiley Interdiscip Rev Dev Biol 2, 47–63.

Wilhelm, L. P., Wendling, C., Vedie, B., Kobayashi, T., Chenard, M. P., Tomasetto, C., Drin, G. and Alpy, F. (2017). STARD3 mediates endoplasmic reticulum-to-endosome cholesterol transport at membrane contact sites. EMBO J 36, 1412–1433.

Xu, J., Liu, H., Lan, Y., Adam, M., Clouthier, D. E., Potter, S. and Jiang, R. (2019). Hedgehog signaling patterns the oral-aboral axis of the mandibular arch. Elife 8.

Yang, X., Li, C., Herrera, P. L. and Deng, C. X. (2002). Generation of Smad4/Dpc4 conditional knockout mice. Genesis 32, 80–81.

Zhang, Y., Liu, T., Meyer, C. A., Eeckhoute, J., Johnson, D. S., Bernstein, B. E., Nusbaum, C., Myers, R. M., Brown, M., Li, W., et al. (2008). Model-based analysis of ChIP-Seq (MACS). Genome Biol 9, R137.

Zhao, J., Shi, W., Chen, H. and Warburton, D. (2000). Smad7 and Smad6 differentially modulate transforming growth factor beta -induced inhibition of embryonic lung morphogenesis. The Journal of biological chemistry 275, 23992–23997.

Zhu, J. and Mackem, S. (2011). Analysis of mutants with altered shh activity and posterior digit loss supports a biphasic model for shh function as a morphogen and mitogen. Dev Dyn 240, 1303–1310.

Zhu, J., Nakamura, E., Nguyen, M. T., Bao, X., Akiyama, H. and Mackem, S. (2008). Uncoupling Sonic hedgehog control of pattern and expansion of the developing limb bud. Dev Cell 14, 624–632.

Zuniga, A. (2015). Next generation limb development and evolution: old questions, new perspectives. Development 142, 3810–3820.

Zuniga, A., Haramis, A.-P. G., McMahon, A. P. and Zeller, R. (1999). Signal relay by BMP antagonism controls the SHH/FGF4 feedback loop in vertebrate limb buds. Nature 401, 598–602.

Zuniga, A. and Zeller, R. (2020). Dynamic and self-regulatory interactions among gene regulatory networks control vertebrate limb bud morphogenesis. Curr Top Dev Biol 139, 61–88.

